# Design of a new effector recognition specificity in a plant NLR immune receptor by molecular engineering of its integrated decoy domain

**DOI:** 10.1101/2021.04.24.441256

**Authors:** Stella Cesari, Yuxuan Xi, Nathalie Declerck, Véronique Chalvon, Léa Mammri, Martine Pugnière, Corinne Henriquet, Karine de Guillen, André Padilla, Thomas Kroj

**Affiliations:** PHIM Plant Health Institute, Univ Montpellier, INRAE, CIRAD, Institut Agro, IRD, Montpellier, France; CBS, Univ Montpellier, CNRS, INSERM, Montpellier, France; IRCM, Institut de Recherche en Cancérologie de Montpellier, INSERM U1194, Université de Montpellier, Institut régional du Cancer de Montpellier, Montpellier, F-34298, France

**Author notes:** **Authors for correspondence:** Thomas Kroj, Stella Cesari.

**Keywords:** NLR specificity, plant immunity, MAX effectors, cell death, rice blast

## Abstract

Plant nucleotide-binding and leucine-rich repeat domain proteins (NLRs) are immune sensors that specifically recognize pathogen effectors and induce immune responses. Designing artificial NLRs with new effector recognition specificities is a promising prospect for sustainable, knowledge-driven crop protection. However, such strategies are hampered by the complexity of NLR function. Here, we tested whether molecular engineering of the integrated decoy domain (ID) of an NLR could extend its recognition spectrum to a new effector. To this aim, we relied on the detailed molecular knowledge of the recognition of distinct *Magnaporthe oryzae* MAX (*Magnaporthe* AVRs and ToxB-like) effectors by the rice NLRs RGA5 and Pikp-1. For both NLRs, effector recognition involves physical binding to their HMA (Heavy Metal-Associated) IDs. However, AVR-PikD, the effector recognized by Pikp-1, binds to a completely different surface of the HMA domain compared to AVR-Pia and AVR1-CO39, recognized by RGA5. By introducing into the HMA domain of RGA5 the residues of the Pikp-1 HMA domain involved in AVR-PikD binding, we created a high-affinity binding surface for this new effector. In the *Nicotiana benthamiana* heterologous system, RGA5 variants carrying this engineered binding surface still recognize AVR-Pia and AVR1-CO39, but also perceive the new ligand, AVR-PikD, resulting in the activation of immune responses. Therefore, our study provides a proof of concept for the design of new effector recognition specificities in NLRs through molecular engineering of IDs. However, it pinpoints significant knowledge gaps that limit the full deployment of this NLR-ID engineering strategy and provides hypotheses for future research on this topic.

## INTRODUCTION

Nucleotide-binding and leucine-rich repeat domain (NLR) receptors are key elements of plant immunity. They detect the activity or the presence of specific pathogen-derived effector proteins that are secreted and translocated inside host cells, and activate defense responses and immunity (Jones & Dangl, 2006). Since they confer resistance to many crop diseases, which represent a major threat to agriculture, NLR-coding genes are widely used in crop breeding programs. Extending the ability of NLRs to recognize a broader range of pathogens is a challenge and represents a major goal for improved disease resistance in crops.

NLR receptors have a conserved architecture comprising a central nucleotide-binding (NB) domain, a C-terminal leucine-rich repeat (LRR) domain and a variable N-terminal signaling domain that is mostly of the coiled-coil (CC) or the Toll-Interleukin1/Receptor (TIR) type (Burdett *et al*., 2019). NLRs recognize specific pathogen effectors through different molecular mechanisms: some interact physically with the recognized effector, while others perceive specific changes induced by an effector on a plant target protein or on a decoy protein that mimics the genuine effector target and thus serves as an effector trap (Cesari, 2018; Kourelis & van der Hoorn, 2018). Many plant NLRs harbor one or multiple non-canonical domains integrated into their structure (Cesari *et al*., 2014; Kroj *et al*., 2016; Sarris *et al*., 2016; Stein *et al*., 2018; Bailey *et al*., 2018; Van de Weyer *et al*., 2019). Some of these integrated domains (ID) were shown to be involved in the specific recognition of effectors and thought to be decoy domains derived from proteins targeted by these effectors (Cesari *et al*., 2013; Maqbool *et al*., 2015; Le Roux *et al*., 2015; Sarris *et al*., 2015; Oikawa *et al*., 2020; Maidment *et al*., 2021). NLR-IDs often cluster genetically and function in combination with a second NLR that acts as a signaling executer for the sensor NLR-ID.

Due to the complexity of NLR function, very few studies have explored the potential of engineering NLR-mediated resistance in plants. Changes in effector recognition specificity between different alleles of the flax NLRs L or P was achieved through domain or residue swaps in the LRR domain (Ellis *et al*., 1999; Dodds *et al*., 2001, 2006; Ravensdale *et al*., 2012). Targeted point mutations or random mutational screens succeeded to extend NLR recognition specificities or increase their activation properties to create sensitized NLRs (Harris *et al*., 2013; Stirnweis *et al*., 2014; Segretin *et al*., 2014). An alternative approach is the engineering of decoy proteins. This was successful in the case of PBS1, which activates the NLR RPS5 upon its cleavage by a bacterial protease effector. The cleavage site of PBS1 was replaced by cleavage sites for other protease effectors from bacteria and viruses resulting in RPS5-dependent resistance to these pathogens (Kim *et al*., 2016; Pottinger *et al*., 2020). Finally, a promising strategy consists in engineering IDs either to extend the recognition spectrum of NLRs or to create new specificities.

Current examples of such engineering mainly focus on improving the ability of a particular NLR to recognize different alleles of a specific effector (De la Concepcion *et al*., 2019). However, there is currently no example of structure-guided engineering of an NLR receptor to confer an entirely different recognition specificity.

In rice, the sensor/executer NLR pair *RGA5/RGA4* confers resistance to *Magnaporthe oryzae* isolates carrying the effector genes *AVR1-CO39* or *AVR-Pia*, while the *Pikp-1* and *Pikp-2* NLR pair recognizes isolates expressing *AVR-PikD* (Ashikawa *et al*., 2008; Okuyama *et al*., 2011; Cesari *et al*., 2013). AVR1-CO39, AVR-Pia, and AVR-PikD are sequence-unrelated, but possess highly similar β-sandwich structures characteristic of the *Magnaporthe* AVRs and ToxB-like (MAX) effector family in plant pathogenic Ascomycete fungi (de Guillen *et al*., 2015; Maqbool *et al*., 2015). Both RGA5 and Pikp-1 contain a heavy metal-associated (HMA) ID that is crucial for specific effector recognition through direct binding (Cesari *et al*., 2013; Maqbool *et al*., 2015). These HMA domains share 54% sequence identity and are located at the C-terminus in RGA5 or between the CC and the NB domains in Pikp-1. Structure–function analyses provided detailed insight into RGA5_HMA/AVR-Pia, RGA5_HMA/AVR1-CO39 and Pikp-1_HMA/AVR-PikD binding and established a causal link between these interactions and effector recognition specificities (Maqbool *et al*., 2015; Ortiz *et al*., 2017; Guo *et al*., 2018).

Remarkably, AVR1-CO39 and AVR-PikD bind different surfaces of the HMA domains (Maqbool *et* al., 2015; Guo *et al*., 2018; De la Concepcion *et al*., 2018), suggesting high plasticity in ID-effector interactions (Varden *et al*., 2019). AVR1-CO39 mainly interacts with the α1 helix and β2 strand of RGA5_HMA whereas AVR-PikD recognition by Pikp-1_HMA involves residue side-chains mainly located in β2, β3, β4 and the terminal K262 residue. Although the 3D structure of the AVR-Pia/RGA5_HMA complex has not been determined yet, gel filtration analyses and 3D modeling suggest that AVR-Pia binds RGA5_HMA through the same interface as AVR1-CO39 (Guo *et al*., 2018; Figure 1). A crystal structure of the AVR-Pia/Pikp-1_HMA complex, showing that AVR-Pia interacts with Pikp-1_HMA through its α1 helix and β2 strand, further supports this hypothesis (Varden *et al*., 2019).

**Figure 1:**
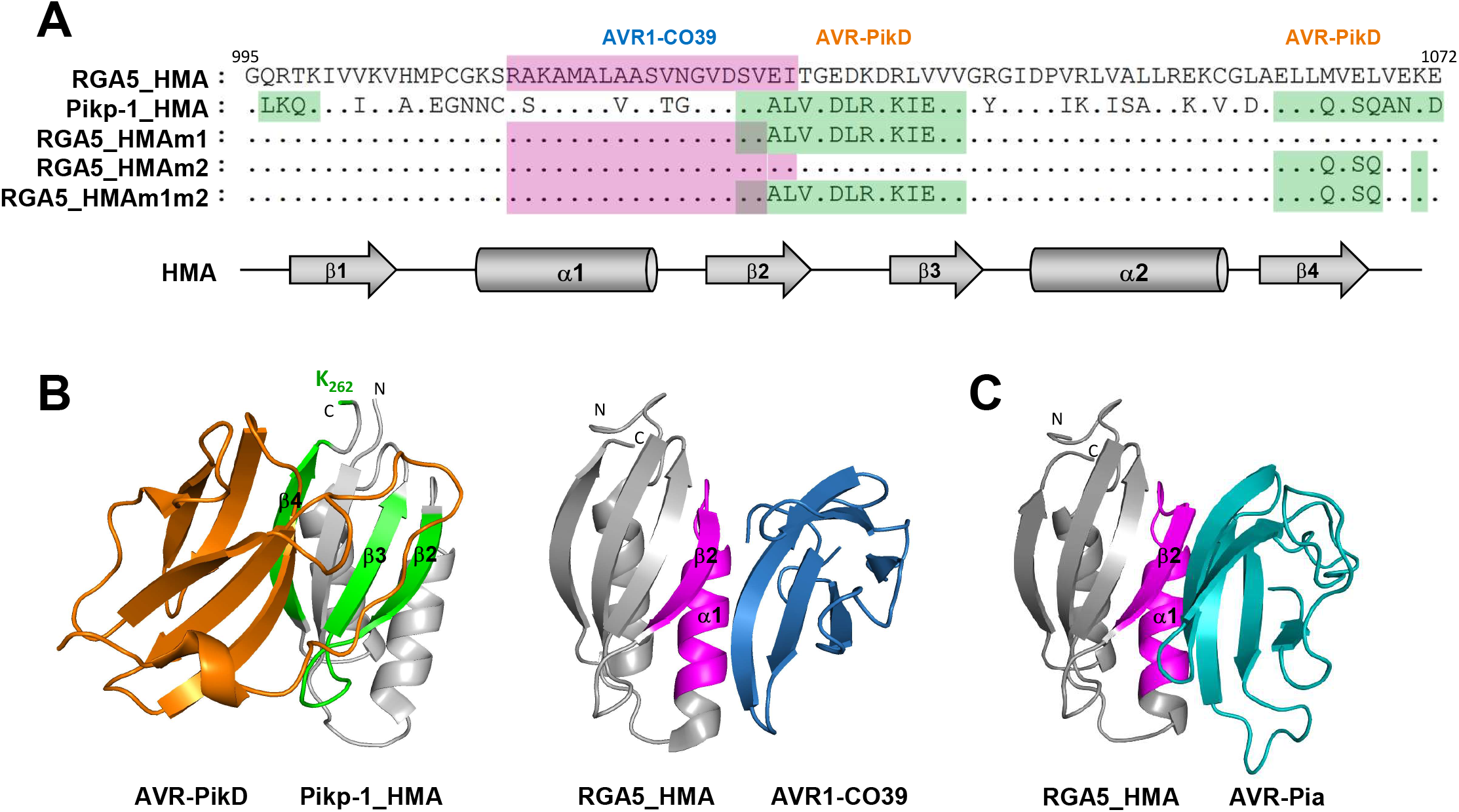
Structure-guided engineering of the HMA domain of RGA5 to introduce the AVR-PikD binding surface. **A)** Sequence alignment of the RGA5 and Pikp-1 HMA domains. The residues constituting the AVR1-CO39 and AVR-PikD binding interfaces are highlighted in pink and green respectively. Residues in strands β2/β3 and β4 targeted respectively by the m1 and m2 mutations are highlighted in green in the different RGA5_HMA variants. Secondary structure elements of the HMA domain are shown below in grey. **B**) Crystal structure of the AVR-PikD/Pikp-1_HMA (left) and AVR1-CO39/RGA5_HMA (right) complexes (PDB_6G10 and PDB_5ZNG respectively). AVR-PikD (orange) binds the HMA domain of Pikp-1 through residues in β2/β3, β4 and K262 (green) and AVR1-CO39 (blue) interacts with the HMA domain of RGA5 through its α1/β2 surface (pink). **C**) 3D model of RGA5_HMA in complex with AVR-Pia (light blue) based on the AVR-Pia/Pikp-1_HMA template structure (PDB_6Q76).

Recent studies have highlighted the potential of ID engineering by altering the recognition specificity of Pik-1 through structure-guided modifications of precise residues within the HMA domain (De la Concepcion *et al*., 2019). This broadened the recognition specificity of the Pik-1 NLR, enabling perception of multiple AVR-Pik alleles.

We hypothesized that by combining the AVR1-CO39 and AVR-PikD binding interfaces in the HMA domain of RGA5, we could generate an RGA5 variant able to bind and recognize both effectors as well as AVR-Pia. We show that introduction of the AVR-PikD binding interface in RGA5_HMA expanded the effector binding capacity of RGA5 *in vivo* and *in vitro*. We report that the engineered RGA5 NLR functions in RGA4 repression and recognizes both AVR-Pia and AVR-PikD in the *Nicotiana benthamiana* heterologous system. However, it does not confer rice resistance to *M. oryzae* isolates expressing AVR-PikD although it still provides resistance to isolates carrying AVR-Pia or AVR1-CO39.

Our study therefore provides a proof of concept that structure-guided engineering is effective to create novel effector-binding interfaces and new recognition specificity for NLR-IDs. However, we also highlight and discuss important constrains and current limitations of the NLR-ID engineering strategy.

## RESULTS

### • Structure-guided engineering of the HMA domain of RGA5

Based on the structure of the Pikp-1_HMA/AVR-PikD complex (Maqbool *et al*., 2015; De la Concepcion *et al*., 2018), we designed mutations in the HMA domain of RGA5 to enable AVR-PikD binding and recognition. The amino acid sequences of the HMA domains of RGA5 and Pikp-1 were aligned (Figure 1A) and the residues of Pikp-1_HMA responsible for AVR-PikD binding were swapped by site-directed mutagenesis into RGA5_HMA generating three RGA5_HMA variants. The m1 mutant harbors nine substituted residues in the β-strands 2 and 3 (E1029A, I1030L, T1031V, E1033D, D1034L, K1035R, R1037K, L1038I and V1039E). The m2 mutant carries three amino acid substitutions in the β4 strand (M1065Q, E1067S and L1068Q) that were defined based on the first published Pikp-1_HMA/AVR-PikD structure (Maqbool *et al*., 2015) and the m1m2 mutant combines the m1 and m2 swaps and thereby harbors almost entirely the AVR-PikD-binding surface of Pikp-1.

The effector-binding surfaces of RGA5_HMA and Pikp-1_HMA are located on opposite sides of the β-sheet forming the conserved core of HMA domains and overlap only slightly on β-strand 2 (Figure 1A and B). Therefore, the m1, m2 and m1m2 mutants remain mostly unchanged for the residues that directly bind AVR1-CO39 (Figure 1A). The structure of the RGA5_HMA/AVR-Pia complex has not be determined yet. However, functional data and the structure of a complex formed between AVR-Pia and Pikp1_HMA suggest a strong overlap between the AVR-Pia and the AVR1-CO39 binding surfaces in RGA5_HMA (Guo *et al*., 2018; Varden *et al*., 2019). Structural modelling of the RGA5_HMA/AVR-Pia complex supports this hypothesis (Figure 1C, Suppl. Table 1). Therefore, the m1, m2 and m1m2 mutations were expected not to interfere with the binding of AVR-Pia to RGA5_HMA.

To address more precisely the potential impact of the m1 and m2 mutations on binding to AVR-Pia, AVR1-CO39 and AVR-PikD, we modelled the structure of the complexes formed between RGA5_HMAm1m2 and all three effectors and calculated corresponding binding energies (Suppl. Figure 1, Suppl. Table 1). This provided similar binding interfaces and energies for the complexes of AVR-Pia and AVR1-CO39 with RGA5_HMA or RGA5_HMAm1m2. Binding parameters of AVR-PikD/RGA5_HMAm1m2 were similar to those in the AVR-PikD/Pikp_HMA complex in both binding surface and binding energy (Suppl. Figure 1, Suppl. Table 1). Detailed *in silico* analysis therefore supported the hypothesis that the RGA5_HMA mutants retain the ability to bind AVR1-CO39 and AVR-Pia and that at least RGA5_HMAm1m2 has high affinity for AVR-PikD.

### • Engineered HMA domains of RGA5 bind AVR-PikD in yeast two-hybrid assays

Using yeast-two-hybrid assays, we found that AVR-PikD interacts with the m1 and m1m2 mutants of both the isolated RGA5_HMA domain (residues 991 to 1072) and a longer C-terminal fragment of RGA5 (residues 883 to 1116) (Figure 2). This RGA5_C-ter fragment includes the sequence downstream of the HMA and part of the linker connecting the LRR and HMA domains and has *in vitro* the same AVR-Pia and AVR1-CO39-binding characteristics as the isolated RGA5_HMA domain (Guo *et al*., 2018). These interactions of AVR-PikD with the m1 and m1m2 mutant constructs were as strong as the one observed with Pikp-1_HMA as shown by yeast growth on stringent selective media supplemented with 10 mM of 3-amino-1,2,4-triazole (3AT). AVR-PikD did not bind RGA5_HMA and, as already described, binds very weakly RGA5_C-ter (Cesari *et al*., 2013). Both RGA5_HMA and RGA5_C-ter m2 mutants also failed to interact with AVR-PikD indicating that changing the corresponding residues in the RGA5_HMA β strand 4 are not sufficient for engineering strong binding. As previously reported, RGA5_C-ter bound strongly to AVR1-CO39 and AVR-Pia (Cesari *et al*., 2013) but, unexpectedly, RGA5_HMA did not. The three RGA5_C-ter mutant variants interacted with AVR-Pia almost at the same level as wild type RGA5_C-ter, but their interaction with AVR1-CO39 was reduced. Indeed, the m1 and m2 mutations slightly decreased binding to AVR1-CO39, while the m1m2 strongly weakened this interaction but did not abolish it (Figure 2, Suppl. Figure 2). Overall, interactions observed for AVR-PikD were stronger than the ones detected for AVR-Pia and AVR1-CO39 as seen on stringent selection conditions (TDO + 10 mM 3AT). We detected all proteins fused to the GAL4 activation domain (AD) or DNA-binding domain (BD) by western blot (Suppl. Figure 3). These results show that modification of the RGA5_HMA surface composed of β strands 2 and 3 is sufficient to confer AVR-PikD-binding and does not or only moderately affect AVR-Pia and AVR1-CO39 binding. Polymorphic residues within the m2 area seem to have limited influence on the binding of AVR-PikD to RGA5_HMA.

**Figure 2:**
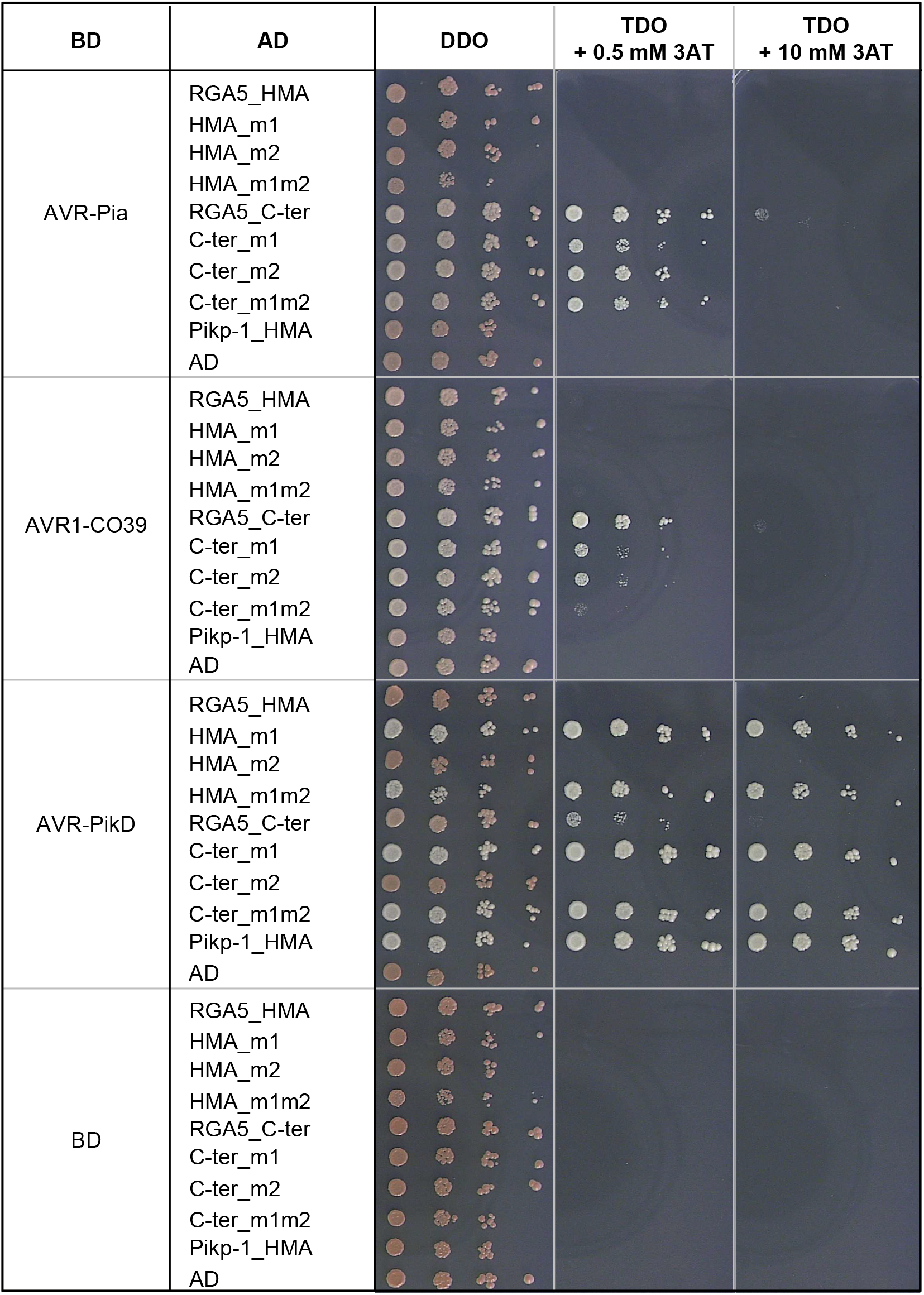
Engineering of the HMA domain of RGA5 enables binding of AVR-PikD in yeast. Interaction of AVR-PikD, AVR1-CO39 and AVR-PikD (without signal peptides and fused to the DNA binding domain (BD) of the GAL4 transcription factor) with the HMA (residues 991 to 1072) and C-terminal (residues 883 to 1116) domains of RGA5 and variants carrying mutations designed to introduce the AVR-PikD-binding surface (fused to GAL4 activation domain (AD)) was assayed by yeast two-hybrid experiments. The HMA domain of Pikp-1 (AD:HMA_Pikp-1) and the AD and BD domains alone were used as controls. Four dilutions of diploid yeast clones (1/1, 1/10, 1/100, 1/1000) were spotted on synthetic TDO medium (-Trp/-Leu/-His) supplemented with 0.5 mM and 10 mM of 3-amino-1,2,4-triazole (3AT) to assay for interactions and on synthetic DDO (-Trp/-Leu) to monitor proper growth. Pictures were taken after 5 days of growth.

### • Engineered RGA5_HMA domains interact strongly with AVR-PikD in vitro

To further characterize these interactions *in vitro*, we performed Surface Plasmon Resonance (SPR) experiments with recombinant HMA domains fused to the Maltose Binding Protein (MBP) and 6xHis-tagged AVR effectors expressed in *Escherichia coli* and purified to homogeneity by affinity chromatography. The MBP-tagged HMA domains were captured on a chip and response units (RU) were measured following the injection of the different AVRs at 1 μM (Figure 3). Comparison of the binding profiles revealed a tight interaction of the AVR-PikD effector with both RGA5_HMAm1 and RGA5_HMAm1m2, similar to that observed with Pikp-1_HMA, whereas poor binding was detected with wild type RGA5_HMA (Figure 3C and D). This was confirmed by performing successive injections of AVR-PikD at increasing concentrations (Suppl. Figure 4). Equilibrium binding constants (K_D_) calculated from these data indicate that the binding affinity of AVR-PikD for both RGA5_HMA mutants is in the nano-molar range as for Pikp1_HMA while its affinity for wild-type RGA5_HMA is in the μ-molar range and thus three to four orders of magnitudes lower (Suppl. Table 2). AVR1-CO39 and AVR-Pia bound equally well to RGA5_HMA mutants and wild type. This binding was rather weak which is consistent with the previously reported μ-molar binding affinity of RGA5_HMA for both ligands (Figure 3A, B and D) (Guo *et al*., 2018). Pikp-1_HMA did not bind AVR1-CO39 but showed, as already described, weak binding to AVR-Pia (Varden *et al*., 2019). The AVR1-CO39_T41G and AVR-Pia_F24S mutants that do not interact with RGA5_HMA and are not recognized by RGA4/RGA5 (Ortiz *et al*., 2017; Guo *et al*., 2018) failed to bind any of the tested HMA domains (Figure 3A and B).

**Figure 3:**
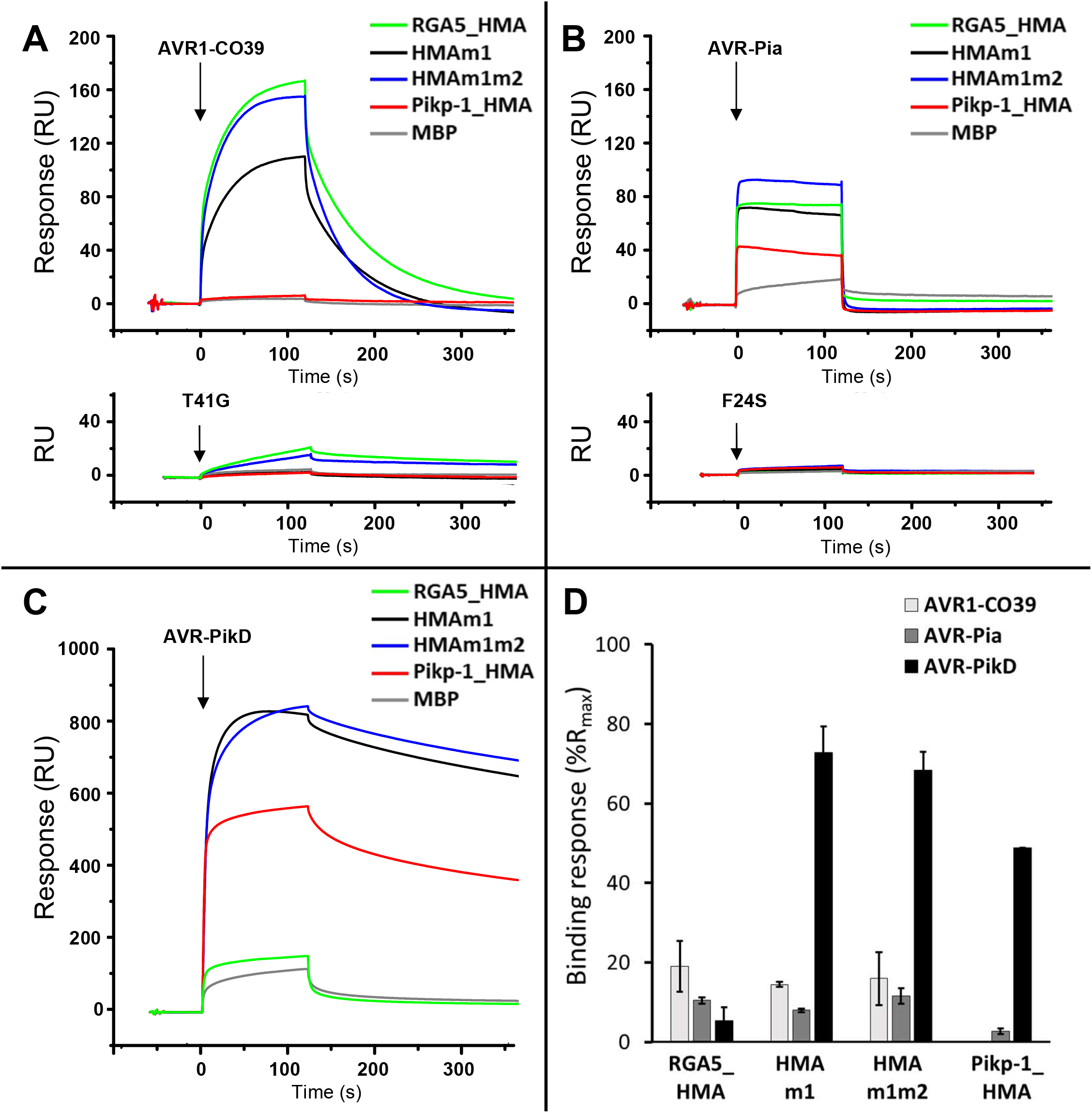
Engineered RGA5_HMAs bind strongly to AVR-PikD *in vitro*. **A-C** AVR1-CO39 (**A**), AVR-Pia (**B**) and AVR-PikD (**C**) were injected (black arrows) at 1 μM for 2 minutes on the different MBP:HMA fusion proteins captured by anti-MBP antibody immobilized on the chip. Superimposed sensorgrams are shown for wild-type RGA5_HMA (green), RGA5_HMAm1 (black), RGA5_HMAm1m2 (blue), Pikp-1_HMA (red), as well as for MBP alone (grey) that serves as negative control. The binding curves obtained with the wild-type and an inactive variant of AVR1-CO39 (**A**) or AVR-Pia (**B**) are shown in the top and lower insets, respectively. **D**) Comparison of the binding response (bound fraction) at 1 μM of AVR effectors, expressed as the percentage of the theoretical maximum response (%Rmax) normalized for the amount of MBP:HMA immobilized on the chip and corrected for the MBP-tag contribution. Bars and error bars represent the mean and average deviation calculated for the %Rmax values estimated from two independent experiments carried out on two different days. Two independently purified protein samples were used to test the binding of the AVR-PikD effector to the different HMAs.

*In vitro* binding is thus consistent with the yeast two-hybrid experiments and supports that engineering the β2-β3 surface of RGA5_HMA creates strong binding to AVR-PikD. Besides, these results demonstrate that the mutations in RGA5_HMA β strands 2, 3 and 4 do not influence binding of AVR1-CO39 or AVR-Pia.

### • Engineered RGA5_HMA domains associates with AVR-PikD in planta

To confirm these interactions *in planta*, we performed co-immunoprecipitation (Co-IP) experiments in *Nicotiana benthamiana* using HA-tagged effectors and YFP-tagged HMA domains. Consistent with yeast two-hybrid and SPR results, AVR-PikD was co-precipitated with RGA5_HMAm1, RGA5_HMAm1m2 and Pikp-1_HMA but did not associate with RGA5_HMA (Figure 4). AVR-Pia specifically associated with the wild type, the m1 and the m1m2 HMA domains of RGA5 but did not co-precipitate with Pikp-1_HMA. We could not test AVR1-CO39:HA because it was not detected in *N. benthamiana* protein extract (Suppl. Figure 5). Interestingly, we observed a correlation between AVR-PikD accumulation in the input and its specific associations with HMA domains suggesting that this effector protein is stabilized upon HMA-binding (Figure 4). As expected, AVR-PikC, a non-recognized allele of AVR-Pik used as negative control was not co-precipitated with any of the HMA domains (Maqbool *et al*., 2015).

**Figure 4:**
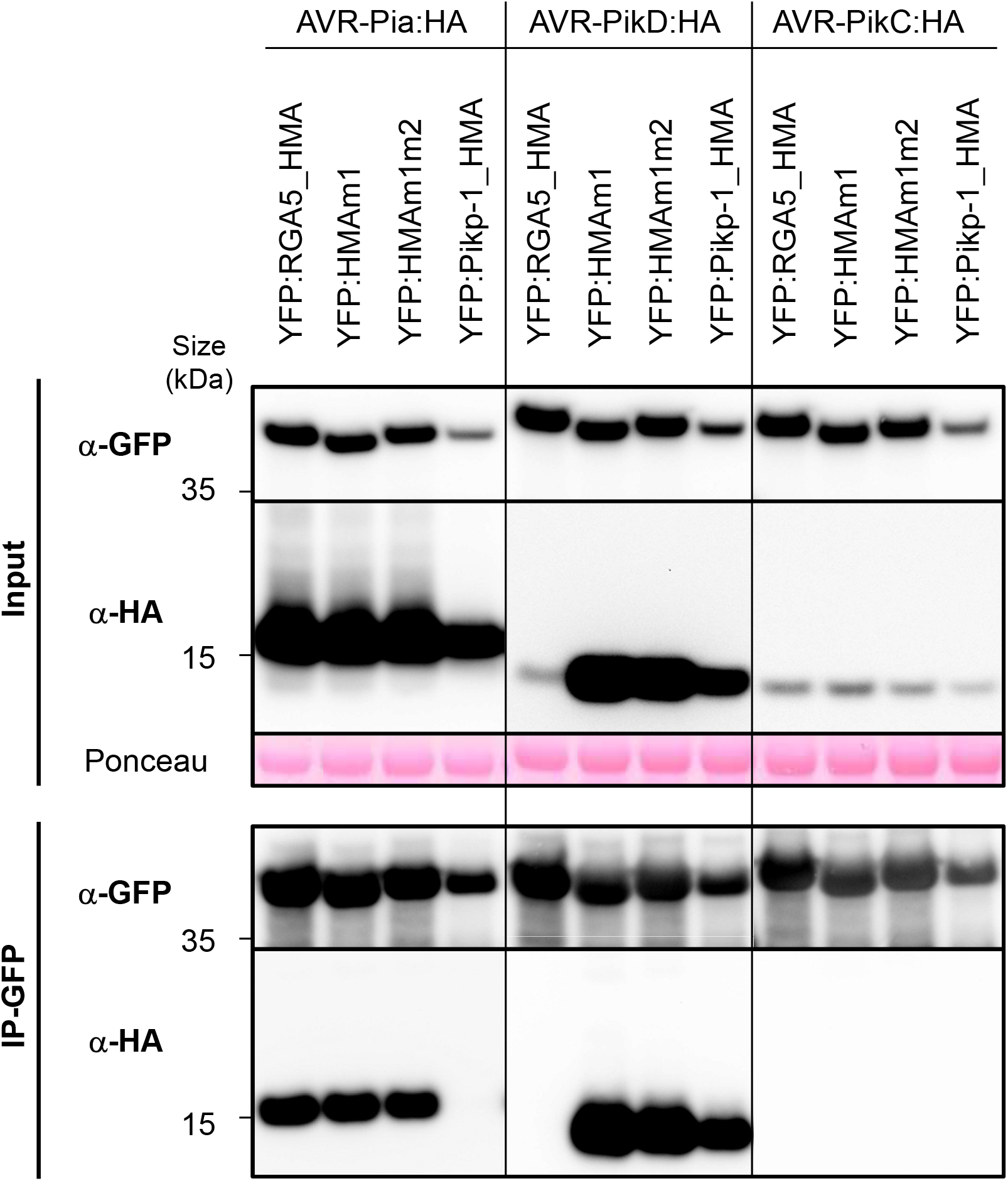
*In planta* association of the engineered HMA domains of RGA5 with MAX effectors. The indicated HMA domains of RGA5 (wild type, m1 and m1m2) and Pikp-1, fused to YFP, were transiently co-expressed in *N. benthamiana* leaves with HA-tagged AVR-Pia, AVR-PikD and AVR-PikC (without signal peptides). Proteins were extracted after 48 hours, separated by gel electrophoresis and tagged proteins were detected in the extract (input) and after immunoprecipitation with anti-GFP beads (IP-GFP, trapping YFP fusions) by immunoblotting with anti-HA (α-HA) and anti-GFP (α-GFP) antibodies. Protein loading in the input is shown by Ponceau staining of the large RuBisCO subunit. The experiment was carried out twice with similar results.

We performed further Co-IP experiments using full-length RGA5 carrying the m1, m2 or m1m2 mutations to test the association of the entire receptors with AVR-Pia and AVR-PikD (Figure 5). HA-tagged effectors and YFP-tagged NLRs were transiently expressed in *N. benthamiana*. Consistent with previous results AVR-PikD was co-precipitated with RGA5m1, RGA5m1m2 and Pikp-1_HMA but did not associate with wild type RGA5 nor with RGA5m2 (Figure 5). AVR-Pia specifically associated with RGA5 wild type, m1, m2 and m1m2 but did not co-precipitate with Pikp-1_HMA. AVR-PikC and PWL2, an unrelated non-MAX effector of *M. oryzae*, did not associate with any of the NLRs. To further exclude that AVR-PikD interacts with other domains of RGA5, we performed Co-IP using a YFP-tagged RGA5 construct deleted for the HMA domain (YFP:RGA5_ΔHMA), and analyzed its association with HA-tagged AVR-PikD alongside HA-tagged AVR-Pia used as a positive control and YFP:Pikp-1_HMA serving as a control for effective and specific co-precipitation of AVR-PikD. We found that YFP:RGA5_ΔHMA co-precipitates AVR-Pia, as previously reported, but not AVR-PikD (Figure 6) (Ortiz *et al*., 2017).

**Figure 5:**
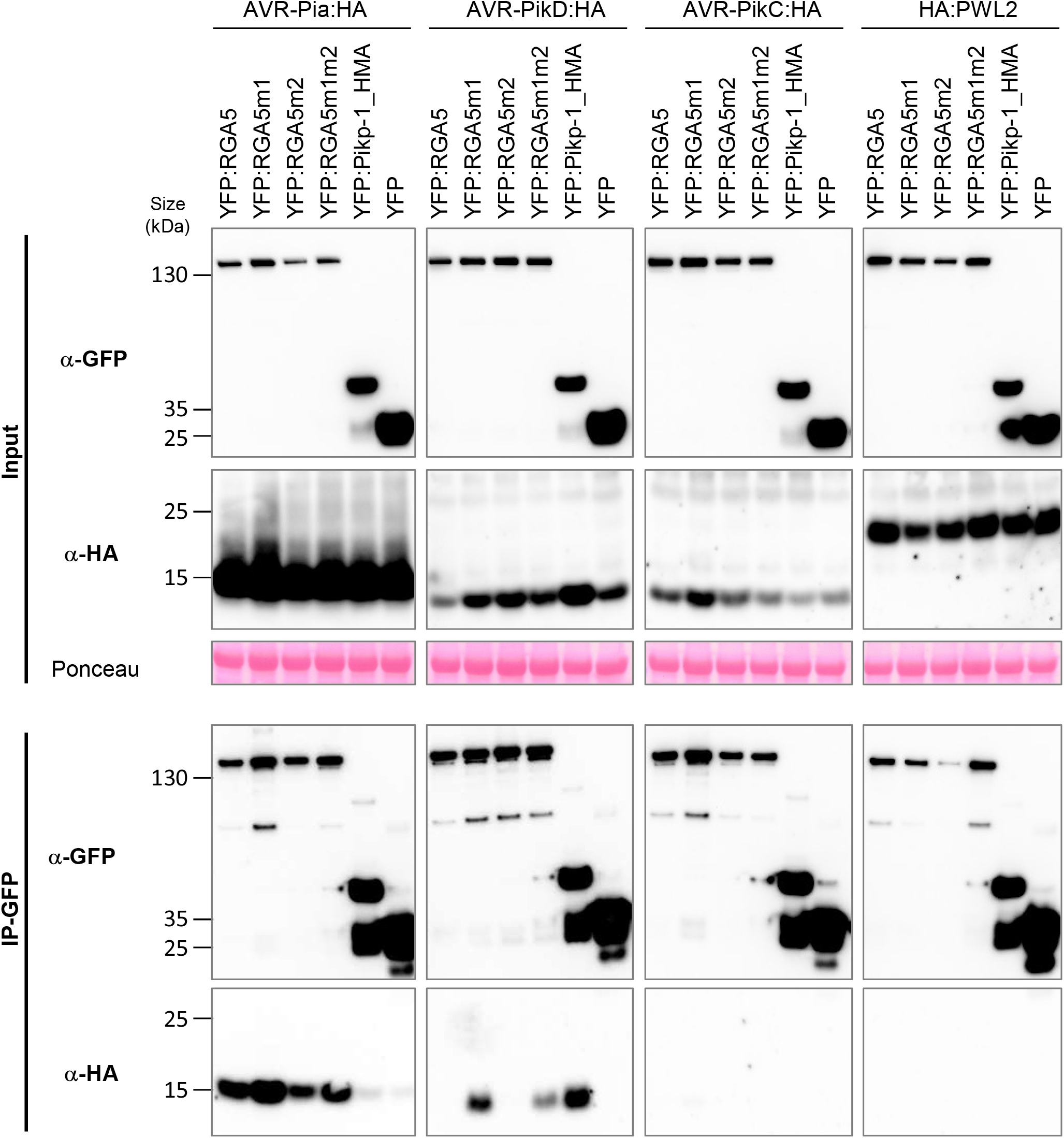
*In planta* association of the engineered full-length RGA5 receptors with MAX effectors. The full-length RGA5 protein and related m1, m2 and m1m2 variants fused to YFP were transiently coexpressed in *N. benthamiana* leaves with the indicated HA-tagged effectors (without signal peptides). The HMA domain of Pikp-1 (fused to YFP) was used as a control as well as YFP alone and the *M. oryzae* effector PWL2. Proteins were extracted 48 hours after infiltration, separated by gel electrophoresis and tagged proteins were detected in the extract (input) and after immunoprecipitation with anti-GFP beads (IP-GFP) by immunoblotting with anti-HA (α-HA) and anti-GFP (α-GFP) antibodies. Ponceau staining shows equal protein loading in the input. The experiment was carried out twice with identical results.

**Figure 6:**
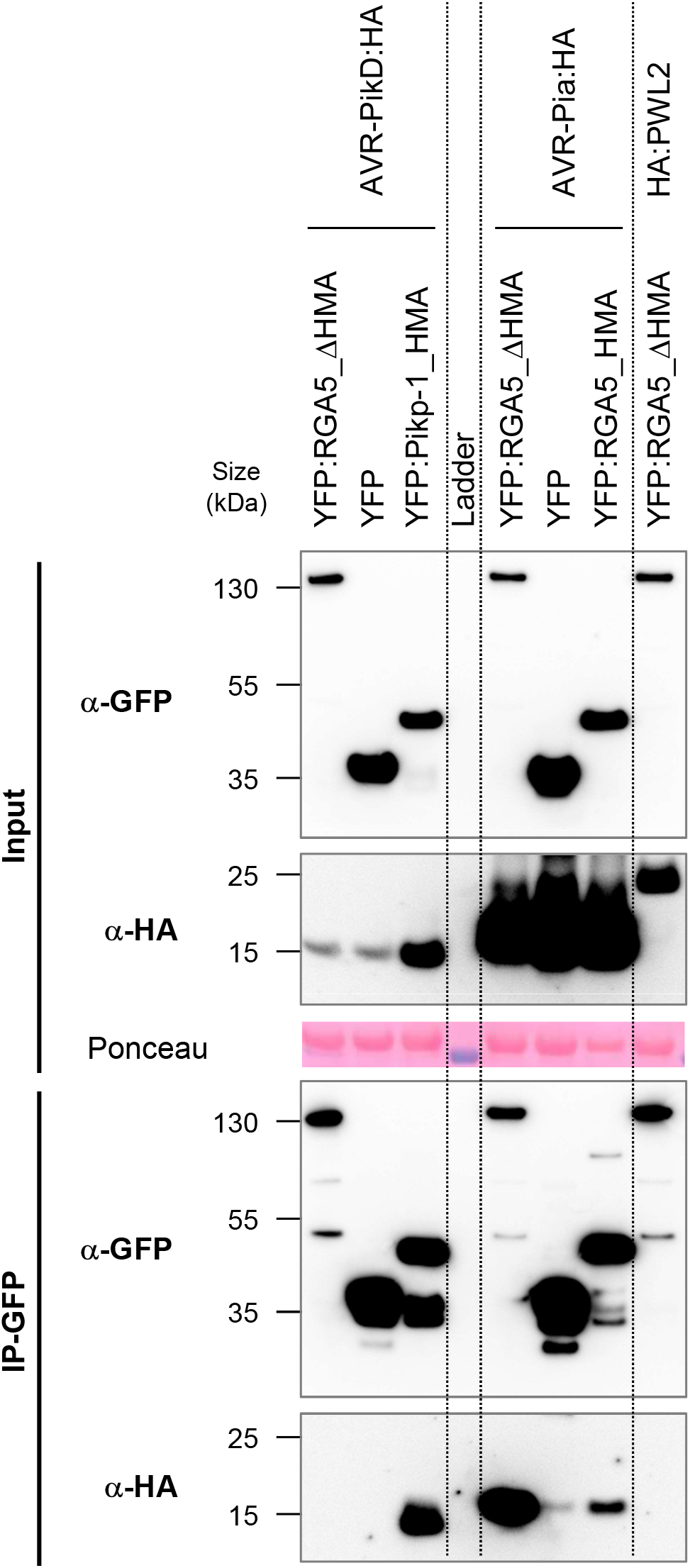
AVR-Pia associates with RGA5_ΔHMA but not AVR-PikD. RGA5 deleted of its C-terminal domain (RGA5_ΔHMA, residues 1 to 996) fused to YFP was transiently co-expressed in *N. benthamiana* leaves with HA-tagged AVR-Pia, AVR-PikD or PWL2 (without signal peptides). Proteins were extracted after 48 hours and tagged-proteins were detected in the extract (input) and after immunoprecipitation with anti-GFP beads (IP-GFP) by immunoblotting with anti-HA (α-HA) and anti-GFP (α-GFP) antibodies. Protein loading in the input is shown by Ponceau staining of the large RuBisCO subunit. The HMA domains of Pikp-1 and RGA5 were used as controls as well as the unrelated PWL2 effector of *M. oryzae*. The experiment was carried out twice with identical results.

Taken together, these experiments indicate that AVR-PikD binds RGA5m1 and m1m2 with high affinity and exclusively through the mutant HMA domains but does not interact with wild-type HMA nor with the rest of the RGA5 protein.

### • The engineered RGA5 variants recognizes both AVR-Pia and AVR-PikD in *Nicotiana benthamiana*

To test whether the m1, m2 and m1m2 mutations enable AVR-PikD recognition by RGA5 in a whole plant context, we performed cell death assays using agro-infiltration of *N. benthamiana* leaves. RGA4 used as a positive control induced cell death when expressed alone but was repressed by RGA5m1m2 as effectively as with wild type RGA5 (Figure 7). This indicates that the m1m2 mutation does not affect the functional interaction between RGA5 and RGA4. Upon co-expression of AVR-PikD and RGA4/RGA5m1m2, a cell death response was induced showing that the m1m2 mutation enables AVR-PikD recognition by RGA5 in the heterologous *N. benthamiana* system (Figure 7). As controls, we observed that AVR-PikD was not recognized by RGA4/RGA5 but induced a strong cell death response upon co-expression with Pikp-1/Pikp-2 (Figure 7, Suppl. Figure 6 and 7). AVR-Pia was specifically recognized by RGA4/RGA5 and RGA4/RGA5m1m2 but not by Pikp-1/Pikp-2 (Figure 7, Suppl. Figure 6).

**Figure 7:**
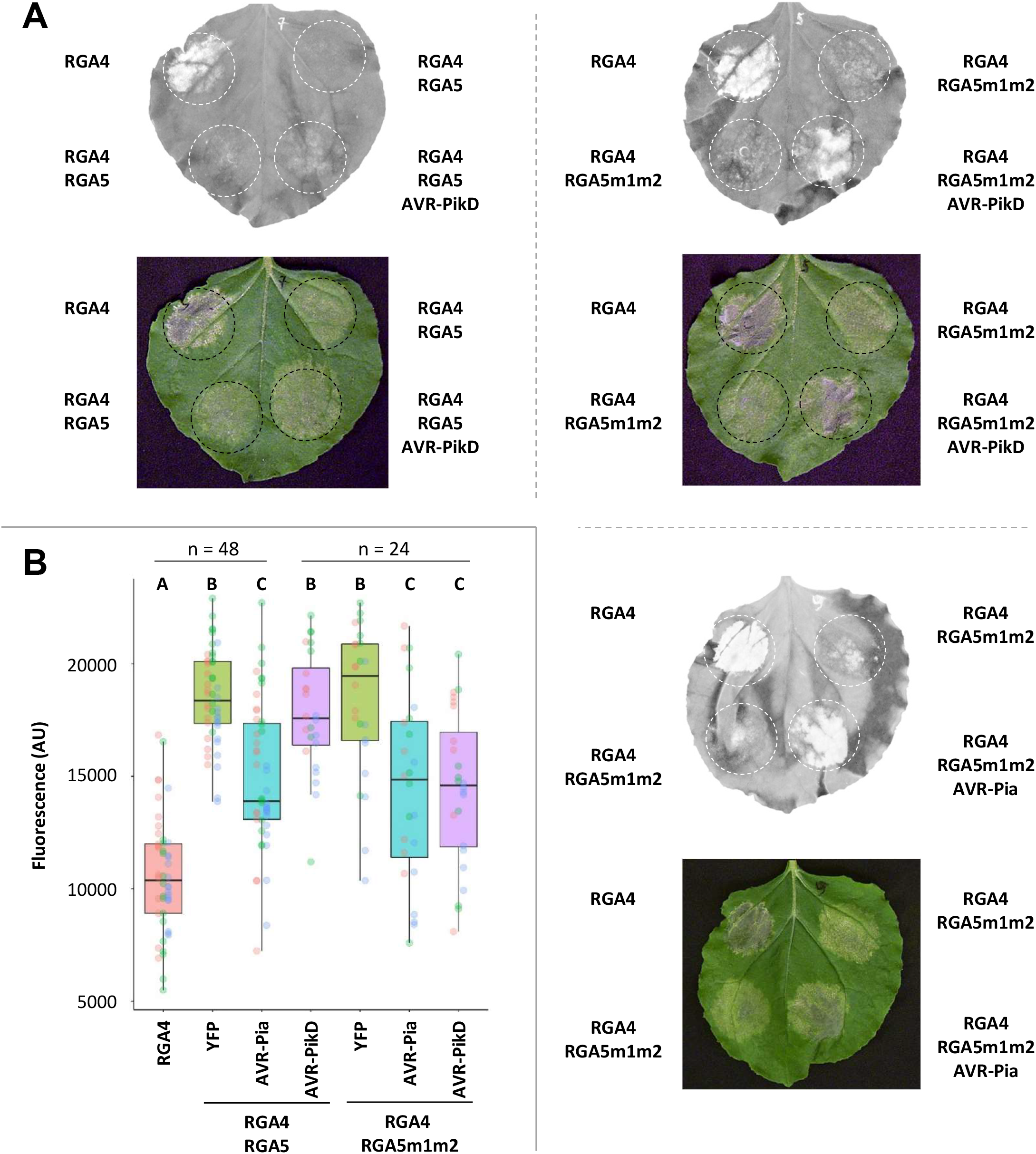
The RGA5m1m2 mutant recognizes AVR-Pia and AVR-PikD in *N. benthamiana*. **A**) The indicated combinations of constructs were transiently expressed in *N. benthamiana* leaves. The AVR-PikD (without signal peptide) and RGA4 constructs carry an HA tag at their C-termini, while RGA5 and RGA5m1m2 are YFP-tagged at their N-termini and AVR-Pia (without signal peptide) is untagged. The auto active RGA4 construct was used as a positive control for cell death induction. Cell death was visualized 5 days after infiltration. Greyscale pictures were taken using a fluorescence scanner with settings allowing visualization of the disappearance of red fluorescence due to cell death (white patches of dead cells). Picture of corresponding leaves are shown below. **B**) Cell death was quantified by measuring fluorescence levels (arbitrary unit, AU) in the infiltrated areas using ImageJ (Xi *et al*., Under review). Resulting data were plotted. The boxes represent the first quartile, median, and third quartile. Difference of fluorescence levels was assessed by an ANOVA followed by a Tukey HSD test. Groups with the same letter (A to C) are not significantly different at level 0.05. For each combination of constructs, all of the measurements are represented as dots with a distinct color (red, green and blue) for each of the three biological replicates.

We also tested AVR-Pia and AVR-PikD recognition by RGA4 and RGA5 carrying only the m1 or m2 surfaces (Suppl. Figure 7). RGA5m1 recognized AVR-PikD but to a lesser extent than RGA5m1m2. The m1 mutation did not affect AVR-Pia recognition. RGA5m2 recognized AVR-Pia but did not trigger cell death upon co-expression with AVR-PikD. All proteins expressed in *N. benthamiana* (besides the untagged AVR-Pia) were detected by western blotting (Figure 5, Suppl. Figure 5).

Consistent with the yeast-two hybrid, Co-IP and SPR data, these results indicate that, in the heterologous *N. benthamiana* system, the engineered RGA5 m1 and m1m2 efficiently bind and recognize both AVR-PikD and AVR-Pia whereas only AVR-Pia is bound and recognized by wild-type RGA5 and RGA5m2.

### • The engineered RGA5 variants do not recognize AVR-PikD but retain AVR-Pia and AVR1-CO39 recognition in transgenic rice plants

To determine whether the m1, m2 and m1m2 mutations of RGA5 confer AVR-PikD recognition in the homologous rice system, we co-transformed the rice cultivar Nipponbare (*pia^-^/pikp^-^*) with constructs carrying under the control of their native promoter the genomic sequences of *RGA4* and *RGA5m1, m2, m1m2* or wild type. Transgenic rice lines transformed with *RGA4* and the *pUbi::GFP* construct were generated as controls. T0 transgenic plants were PCR-genotyped for the presence of *RGA4* and *RGA5* and all PCR products corresponding to m1, m2 and m1m2 transgenic lines were sequenced to ensure that they contain the appropriate mutations (Suppl. Table 3). This identified 14 independent T0 transgenic lines successfully transformed with *RGA4/RGA5*, 6 with *RGA4/RGA5m1*, 5 with *RGA4/RGA5m2*, 6 with *RGA4/RGA5m1m2* and 14 with *RGA4/GFP* (Suppl. Table 3).

When possible, thalli from individual T0 plants were split and inoculated with the *M. oryzae* isolate Guy11 transformed with either the empty vector (Guy11_EV) or with *AVR-Pia* (Guy11_*AVR-Pia*), or with the wild type strain JP10 naturally carrying *AVR-PikD* and lacking *AVR-Pia*. As expected, all rice transgenic lines inoculated with Guy11_EV showed susceptibility (Suppl. Table 3, Suppl. Figure 8). Upon inoculation with Guy11_*AVR-Pia*, 6/8 *RGA4/RGA5*, 1/2 *RGA4/RGA5m1*, 3/3 *RGA4/RGA5m2* and 3/5 *RGA4/RGA5m1m2* T0 transgenic lines showed resistance, while all those inoculated with JP10 developed disease phenotypes. The K60 rice cultivar carrying the *Pikp-1/Pikp-2 NLR* pair and diagnostic for Pikp resistance was resistant to JP10, indicating proper AVR-PikD recognition (Kiyosawa, 1984; Kanzaki *et al*., 2012). These results indicate that the engineered RGA5m1, m2 and m1m2 variants all recognize AVR-Pia but not AVR-PikD in the homologous rice/*M. oryzae* system.

To backup this analysis, which was performed on T0 transgenic plants that are highly heterogeneous and stressed due to regeneration from *in vitro* culture, inoculation experiments were also carried-out on T1 plants (Suppl. Table 3, Figure 8). For this, we used a transgenic Guy11 isolate transformed with *AVR-PikD* instead of JP10 to ensure homogeneous fungal background. We also included the Guy11_AVR1-CO39 transgenic isolate. Based on seed availability, 3 independent rice transgenic lines were selected for *RGA4+RGA5* and *RGA4+RGA5m2*, 2 for *RGA4+RGA5m1* and 1 for *RGA4+RGA5m1m2*. Consistent with the results obtained in the T0 experiment, AVR-Pia was recognized by RGA5 and its variants, but not AVR-PikD. In addition, like RGA5 wild type, RGA5m1, m2 and m1m2 also recognized AVR1-CO39.

**Figure 8:**
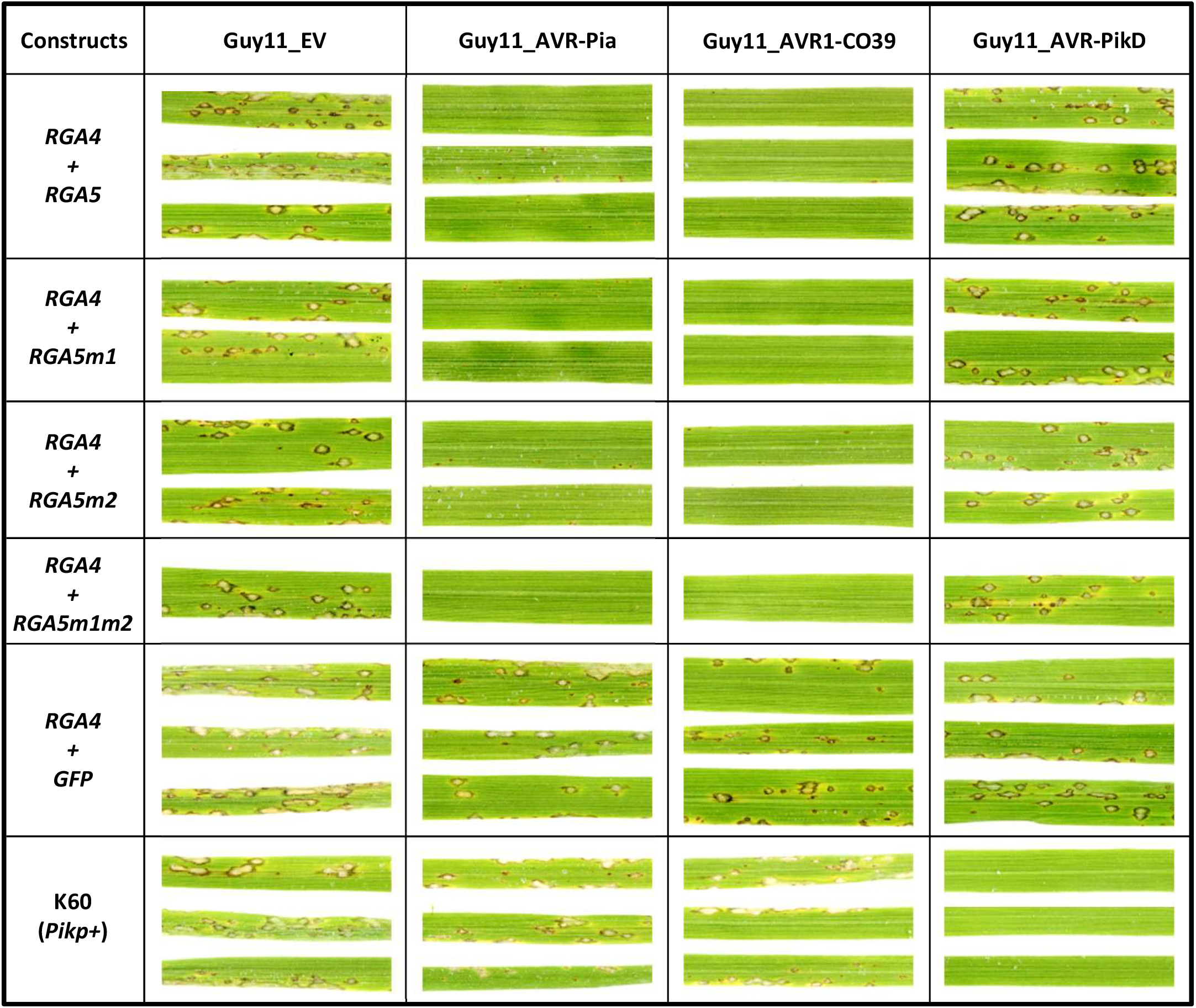
The RGA5 m1, m2 and m1m2 mutants recognize AVR-Pia but not AVR-PikD in rice. The rice cultivar Nipponbare was co-transformed with a genomic construct for *RGA4* and a genomic construct for *RGA5, RGA5m1, RGA5m2* or *RGA5m1m2*. A transgenic line carrying *RGA4* and *GFP* was also generated as a control. T1 plants of the transgenic lines were spray inoculated with the transgenic strains Guy11-AVR-Pia, Guy11-AVR1-CO39, Guy11-AVR-PikD or Guy11-EV. The rice cultivar K60 carrying the *Pikp* resistance was used as a control for AVR-PikD specific recognition. Pictures show representative symptoms at 7 days after inoculation. Individual leaves indicate independent T1 transgenic lines. Similar results were obtained in two independent inoculation experiments performed on T0 (Suppl. Figure 8) and T2 (Suppl. Figure 9) plants.

To identify a potential partial resistance to AVR-PikD in the transgenic lines, we conducted inoculation experiments using T2 plants (Suppl. Table 3, Suppl. Figure 9) and precisely measured lesion size using the computational image analysis package Leaftool (https://github.com/sravel/LeAFtool). These experiments show that in the transgenic RGA5m1, m2 and m1m2 plants disease lesions caused by Guy11_AVR-PikD are not reduced in size compared to the control rice lines or to infection with the Guy11_EV isolate. However, AVR-Pia-induced resistance manifested by small hypersensitive response-type lesions equally induced in all RGA5 lines (Suppl. Figure 9). This further confirms that, in rice, AVR-PikD is not recognized by the engineered RGA5 variants.

## DISCUSSION

NLR immune receptors provide plants with efficient protection from biotrophic pathogens and the corresponding genes are therefore critical for breeding disease resistant crops. However, not all NLR resistance genes are useful for crop protection since their spectrum can be extremely limited due to the polymorphism of effector-coding genes. In certain cases, no NLR inducing broad-spectrum resistance is present in crop germplasm. A promising prospect to overcome these limitations is creating NLRs with specific recognition specificities by molecular engineering (Cesari, 2018). NLR-IDs have been recognized as prime candidates for such resistance engineering due to their modular structure and the well-established role of decoy IDs in effector recognition (Cesari *et al*., 2013; Le Roux *et al*., 2015; Sarris *et al*., 2015).

In the RGA4/RGA5 and Pik1/Pik2 model systems, detailed structure-function analysis deciphered at atomic scale how IDs contribute to the recognition of fungal effector proteins (Maqbool *et al*., 2015; Ortiz *et al*., 2017; Guo *et al*., 2018; De la Concepcion *et al*., 2018). These studies established that small HMA proteins have been recruited repeatedly and independently to serve as decoy domains that physically bind effectors like molecular traps (Oikawa *et al*., 2020; Maidment *et al*., 2021). These breakthroughs pave the way towards rational structure-guided design of effector-binding domains in these NLRs. First proof of concept studies in the Pik-1/Pik-2 system demonstrated how structure-guided ID engineering enabled the recognition of different alleles of the same effector (De la Concepcion *et al*., 2019). However, the creation of entirely different specificities by ID engineering has not been reported yet.

### • Rational structure-guided design of an HMA domain confers novel effector binding

In this study, we successfully generated a new effector recognition specificity for the RGA5 immune receptor by creating a novel binding surface in its HMA ID. For this, we swapped a limited number of residues from the Pikp-1_HMA into the RGA5_HMA domain. The identification of critical residues to be mutated relied on structural knowledge obtained at high resolution for these HMA domains in complex with different AVRs (Maqbool *et al*., 2015; Guo *et al*., 2018; De la Concepcion *et al*., 2018). The resulting mutants, RGA5_HMAm1 and RGA5_HMAm1m2, were assessed *in silico* for effector binding by modelling of relevant effector/HMA complexes. These models predicted that both RGA5_HMA mutants could form complexes with AVR1-CO39, AVR-Pia and AVR-PikD and that binding affinities in these complexes would be similar to those in the respective complexes formed by wild type RGA5_HMA and Pikp1_HMA. *In vivo* and *in vitro* experiments confirmed these predictions and demonstrated that RGA5_HMAm1 and RGA5_HMAm1m2 bind AVR-PikD with high affinity in addition to retaining moderate binding affinity to AVR-Pia and AVR1-CO39.

The K_D_ values retrieved from our SPR experiments indicated nano-molar affinity (ranging from 0.5 to 1.6 nM, Suppl. Table S2) for the complexes formed between AVR-PikD and the engineered RGA5_HMAs, similar to that estimated for the complex formed with the wild type Pikp-1_HMA. This is consistent with the K_D_ value of 6 nM formerly determined by SPR for AVR-PikD/Pikp-1_HMA interaction using different constructs and experimental settings (De la Concepcion *et al*., 2018). Therefore, our results with the RGA5_HMA mutants further illustrate the previously established critical role of the HMA β2 and β3 strands in AVR-PikD binding and recognition (Maqbool *et al*., 2015; De la Concepcion *et al*., 2018). The fact that m2 mutations have a low impact on AVR-PikD binding to RGA5_HMA is also consistent with the limited contribution of the β4 strand of Pikp-1_HMA to AVR-PikD specific recognition (De la Concepcion *et al*., 2019). However, this β4 strand is important for the specificity of recognition of other AVR-Pik alleles by Pik-1 alleles (De la Concepcion *et al*., 2018, 2019; Varden *et al*., 2019). The affinity of AVR-Pia or AVR1-CO39 to both wild type RGA5_HMA and RGA5_HMAm1m2 was in the μ-molar range as measured here by SPR, matching the K_D_ values of AVR1-CO39/RGA5_HMA (5 μM) and AVR-Pia/RGA5_HMA (7.8 μM) previously determined by ITC measurements (Ortiz *et al*., 2017; Guo *et al*., 2018). This weak impact of the m1 and m2 mutations on AVR-Pia and AVR1-CO39 binding further confirms the current structural model for the detection of these effectors and nicely illustrates the fact that different MAX effectors can bind two different interfaces in the HMA domain.

### • Binding to the ID is sufficient for recognition in the heterologous *N. benthamiana* system but not in the homologous rice system

Previous work established the central role of HMA binding in the detection of pathogen effectors by the RGA5 and Pikp-1 immune receptor (Maqbool *et al*., 2015; Ortiz *et al*., 2017; Guo *et al*., 2018). Indeed, effector mutants that are strongly impaired in binding to the HMA lose their avirulence activity and do no longer trigger immunity in resistant rice cultivars. The RGA5m1 and m1m2 mutants created in this study gain high affinity binding to AVR-PikD and activate cell death immune responses when coexpressed with this effector in the heterologous *N. benthamiana* model system. This indicates that the new-to-nature receptors harboring the engineered HMA domains are capable of forming functional assemblies responding to AVR-PikD in addition to the naturally recognized effectors AVR-Pia and AVR1-CO39. However, the cell death response is similar or slightly weaker than with AVR-Pia although the binding of RGA5_HMAm1m2 to AVR-PikD detected in yeast or by SPR is much stronger than to AVR-Pia and AVR1-CO39. Similar discrepancies between the gain-of-binding observed *in vitro* and the eliciting of the cell-death response in *N. benthamiana* has also been reported for a Pikp-1 variant with an engineered HMA domain (De la Concepcion *et al*., 2019). Therefore, for RGA5 as for Pikp-1, the effector/HMA binding affinity does not necessarily correlate with the strength of the induced immune response. This suggests that additional factors may influence effector recognition by these NLRs.

When transformed into rice, RGA5m1 and RGA5m1m2 do not confer resistance to *M. oryzae* isolates carrying *AVR-PikD* although they are as active as wild-type RGA5 in recognizing *AVR1-CO39* and *AVR-Pia* isolates. This indicates that in the homologous rice system the binding of an effector to the HMA domain even with high affinity is not sufficient to activate the RGA4/RGA5 complex and to trigger immune responses. Two main hypothesis can explain this result and the missing correlation between HMA ID binding affinity and response in *N. benthamiana* cell death assays. Effectors may have to bind at the proper side of the RGA5_HMA domain to trigger activation of RGA4/RGA5. This may be due to steric or spatial constraints in the receptor complex or because effectors have to disrupt intramolecular interactions mediated by the AVR-Pia and AVR1-CO39-binding surface of RGA5_HMA. The AVR-Pia and AVR1-CO39-binding surface of RGA5_HMA could also bind rice proteins not present in *N. benthamiana* that have to be replaced by the effectors for proper activation of the immune receptor complex. Alternatively, HMA-binding alone may not be sufficient for efficient receptor activation because additional interactions with RGA5 outside the HMA domain are required for full receptor activation. Support for this hypothesis comes from the finding that AVR-Pia associates with RGA5_ΔHMA in Co-IP experiments while AVR-PikD does not. Such additional interactions between the effector and RGA5 outside of the HMA domain should not only stabilize effector binding and significantly increase overall binding affinity, but may directly induce conformational changes in RGA5 required for efficient RGA4/RGA5 receptor complex activation (Ortiz *et al*., 2017). Additional support for this hypothesis comes from the previous investigation of the recognition of AVR-Pia and AVR1-CO39 mutants by RGA4/RGA5 that showed high resilience to reduction of HMA-effector affinity (Ortiz *et al*., 2017; Guo *et al*., 2018). Indeed, effector mutants with strongly reduced HMA affinity were still recognized and only complete loss of HMA-binding resulted in loss of avirulence activity. Interestingly, the recently reported structures of two different resistosomes composed of the TIR-NLRs ROQ1 or RPP1 in complex with their matching effectors (XopQ and ATR1 respectively) show examples where effectors recognition relies on direct binding to multiple sites and domains of the NLRs (Ma *et al*., 2020; Martin *et al*., 2020). In both cases, the effectors do not only bind an extended surface of the LRR domain but also the post-LRR domain that adopts a jelly roll/Ig-like fold, which occurs specifically and with high frequency in TNLs and is important for receptor function (Van Ghelder & Esmenjaud, 2016; Saucet *et al*., 2021).

The recognition of AVR-PikD by RGA5m1 and m1m2 in *N. benthamiana*, despite a potentially incomplete receptor binding, may be due to the strong overexpression of effectors and receptors in this system. The resulting high cellular levels of receptors and effectors may overcome sub-optimal conditions and enable to reach the threshold that is required for activated receptor complexes triggering immune responses and cell death. Similar discrepancies between effector recognition in *N. benthamiana* cell death assays and pathogen resistance in the homologous system has been reported in a study that aimed to engineer the NLR R3a from potato for recognition of additional alleles of the effector AVR3a from *Phytophthora infestans* (Segretin *et al*., 2014).

### • Perspectives for future attempts of rational NLR design

While NLRs from many different clades can carry IDs, the frequency of NLR-IDs in most NLR clades is low. In cereals, only 3 NLR clades named major integration clades (MIC) are strongly enriched for NLR-IDs (Bailey *et al*., 2018). In MIC2 and MIC3, IDs are conserved (DDE superfamily endonuclease and the BED-type zinc finger domains, respectively), indicating that they originate from unique ancient integration events. Only MIC1 to which belongs RGA5 but not Pikp-1 harbors highly diverse IDs that correspond to many different types of protein domains. Comparison of orthologous MIC1 NLRs indicates frequent and ongoing exchange of IDs (Cesari *et al*., 2014; Bailey *et al*., 2018). Since MIC1 NLRs can accommodate such a huge diversity of IDs and since swap-in of novel IDs seems a common mechanisms and a frequent event in their evolution, they appear particularly suited “chassis” for generating “à la carte” novel effector-recognition specificities by molecular engineering and ID manipulation. Until recently, such engineering was hampered by the lack of knowledge regarding the molecular mechanisms of NLR receptor action, both in the specific recognition of effectors and in their underlying activation. Major breakthroughs in the determination of the 3D structure of effectors and NLR receptors are lifting these barriers and pave the way towards structure-guided engineering of NLR receptors.

Based on structural knowledge on effector-ID interactions, we succeeded in generating artificial NLR receptors that bind a novel effector through a previously not-existing binding interface and have a novel recognition specificity in heterologous *N. benthamiana* cell death assays. However, the finding that these engineered NLRs do not provide a novel resistance specificity in the homologous rice system illustrates our limited understanding of the action of IDs in full-length receptors and of the interactions in NLR pairs at the molecular level. In particular, the role of the binding of effectors to RGA5 outside the ID in effector recognition and RGA4/RGA5 activation is not established. Mapping these interactions and determining their importance by mutant studies is therefore crucial to evaluate if and how they should be considered when novel NLR-ID recognition specificities are generated with MIC1 NLRs.

In addition, it is critical to understand more precisely how the binding of effectors to RGA5 de-represses the RGA4/RGA5 complex. The recent description of the molecular and structural mechanism of effector recognition by the Arabidopsis CNL ZAR1 provides a blueprint for the understanding of CNLs that act alone and rely on decoy or guard proteins for effector recognition (Wang *et al*., 2019b, a). Similarly, detailed studies are required to provide a molecular and structural framework for MIC1 NLR action and their interaction with RGA4-like executor NLRs. It is tempting to speculate that RGA4/RGA5 may form upon activation a similar multimeric resistosome complex as ZAR1/RKS1 (Adachi *et al*., 2019) and form, at the inactive state, a repressed heterodimeric complex. However, to address this hypothesis and to decipher the intra and intermolecular interactions in these complexes and the conformational changes mediating their transitions will require an integrated multidisciplinary approach combining structure biology, *in vitro* biochemistry and functional plant genetics. In addition, it would be highly instrumental to dispose of additional MIC1 NLR model systems where the effectors and their interaction with the ID of the RGA5-like sensor is known to decipher the communalities and the specificities in different MIC1 NLR pairs. In this sense, this study not only illustrates the potential that NLR-IDs provide for creating almost unlimited number of pathogen recognition specificities but also the tremendous challenges that remain to be addressed before this ambitious goal can be reached.

## METHODS

### • Protein expression and purification

Details on the plasmid constructs and synthetic genes used in the present study are given in Suppl. Table 5 and Suppl. Text 1. Wild-type and mutant AVR1-CO39 and AVR-Pia proteins, deleted for their endogenous secretion signal, were expressed and purified as His-tagged proteins from *E. coli* periplasm as previously described (de Guillen *et al*., 2015). For AVR-PikD, the His-tagged protein was expressed in *E. coli* BL21 (DE3) from the pET derived plasmid pdbccdb_3C_His (kindly provided by F. Allemand) and purified under denaturating condition by affinity chromatography (50 mM Tris, pH 8.5, 300 mM NaCl, 1 mM DTT, 8 M urea), refolded by over-night dialysis and further purified by gel filtration in 20 mM Tris, pH 8.5, 150 mM NaCl, 1 mM DTT. Wild-type and mutant HMA domains from RGA5 and Pikp-1 fused to the maltose binding protein (MBP) were expressed from a pdbccdb_3C_His_MBP construct and purified on His-trap column followed by over-night dialysis in 20mM Tris pH 7.5, 150mM NaCl, 1mM DTT. All protein samples were stored at −20°C.

### • SPR

SPR experiments were performed at 25°C on a Biacore T200 apparatus (GE Healthcare) in running buffer (20 mM Tris, pH 8, 150 mM NaCl, 1 mM DTT) supplemented with 0.05% of P20 surfactant (GE Healthcare). Anti-MBP murin monoclonal antibody (Biolabs) was immobilized (around 11000 responsive units (RU)) on CM5S dextran sensor chip (Cytiva) by amine coupling according to the manufacturer’s instructions. The MBP protein alone or fused to the wild-type or mutant HMA domain from RGA5 or Pikp-1 was then injected at 300 nM on the sensor chip, leading to a capture level of 3000-4000 RU. For comparative binding experiments, the concentrated AVR effectors were diluted at 1 μM in running buffer and injected at 30 μL/min during 120 sec followed by a dissociation phase in flow buffer. Cycle was ended by injecting 7 μL of regeneration buffer (10 mM glycine-HCL, pH 2). All curves were analyzed using Biacore T200 BiaEvaluation software 3.0 (GE Healthcare), after double–referencing subtraction. Binding levels (bound fraction) were compared by calculating the fraction of immobilized MBP:HMA with bound AVR at the end of injection, expressed as a percentage of the theoretical maximum response (%Rmax) and corrected for the contribution of the MBP tag by subtracting the bound fraction calculated for the MBP protein alone. For kinetic titration experiments (single-cycle kinetics), the analyte (AVR-PikD) was dialyzed in running buffer (20 mM Tris, pH 8, 150 mM NaCl, 0.5 mM DTT) and increasing concentrations were injected successively on captured MBP:HMAs (about 4000 RU) at a flow rate of 30 μL/min for 60 sec followed by a dissociation phase of 600 sec after the final injection. Data analysis and determination of binding parameters were performed with BiaEvaluation using steady state or heterogeneous binding fitting models to obtain the best fitting().

### • Growth of plants and fungi and infection assays

Rice plants (*Oryza sativa* L.) were grown as described (Faivre-Rampant *et al*., 2008). *Nicotiana benthamiana* plants were grown in a growth chamber at 22°C with a 16 hours light period. *M. oryzae* isolates and transgenic strains were grown as described (Berruyer *et al*., 2003). For the determination of interaction phenotypes and gene expression, a suspension of fungal conidiospores (50000 spores.ml^-1^) was spray-inoculated on the leaves of 3-week-old rice plants. Rice leaves were collected and scanned 7 days after inoculation.

### • Constructs for yeast two-hybrid, Co-IP and rice transformation

PCR products used for cloning were generated using Phusion High-Fidelity DNA Polymerase (Thermo Fisher) using primers listed in Suppl. Table 4. Details of constructs are given in Suppl. Table 5. Briefly, all ENTRY vectors used for LR cloning were obtained either by gateway BP cloning (Life Technologies) into the pDONR207 vector or through site-directed mutagenesis (Quikchange lightning technology, Agilent technologies) using an ENTRY clone as template. Plasmids used for yeast two-hybrid or Co-IP were generated by gateway LR cloning (Life Technologies) using the ENTRY vectors described above and appropriate destination vectors listed in Suppl. Table 5. Plasmids used for rice transformation were created by site-directed mutagenesis (Quikchange lightning technology, Agilent technologies) to introduce point mutations in the genomic sequence of RGA5 already cloned in pAHC17. The resulting constructs were digested using the HindIII and BamHI restriction enzymes (BioLabs) and cloned in the pCambia2300 vector.

### • Yeast two-hybrid analysis

Yeast two-hybrid assays were performed as described (Cesari *et al*., 2013) using the Matchmaker Gold Yeast Two-hybrid system (Clontech).

### • Transient protein expression in *N. benthamiana*

Agro-infiltration in *N. benthamiana* were performed as described (Cesari *et al*., 2013). Leaves were harvested, scanned and analyzed as described by Xi et al. (Xi *et al*., Under review). Boxplots were generated using R v4.0.2 and the package tidyverse (Wickham *et al*., 2019). Difference of red fluorescence induced by the various agro-infiltrated constructs was assessed either by a one-way ANOVA followed by a Tukey HSD test or by a Kruskal Wallis test followed by a Dunn test.

### • Transgenic rice lines

*pRGA5:RGA4, pRGA5:RGA5, pRGA5:RGA5m1, pRGA5:RGA5m2, pRGA5:RGA5m1m2* and *pUBI:GFP* were used for *Agrobacterium tumefaciens*–mediated transformation (strain EHa105) of wild-type Nipponbare rice (Toki *et al*., 2006). Infected calli were selected on medium containing 200 mg.L^-1^ geneticin and 50 mg.L^-1^ hygromycin. Resistant calli were transferred to regeneration medium. T0 plants were used for *M. oryzae* inoculation assays 3 weeks after transfer to soil. For this, regenerated plants with at least three tillers were split in several plantlets and replanted in soil in independent pots. The presence of the transgenes in T0 plants was verified by PCR. Sequencing was used to confirm the identity of the m1, m2 and m1m2 constructs.

### • Protein extraction immunoblot and co-immunoprecipitation

Protein extraction from *N. benthamiana* leaves and co-IP experiments were performed as described previously (Guo *et al*., 2018) with the following modifications: for Co-IP assays using full-length RGA5 (and its variants), a high stringency buffer was used for the extraction (50 mM Tris HCl pH7.5, 150 mM NaCl, 1 mM EDTA, 1% NP40, 0.1% SDS, 0.5% Deoxycholate, 10 mM DTT, 0.5% PVPP, 1% protease Sigma inhibitor and 1 tablet of Roche protease inhibitor for 50 ml of buffer) and the wash steps (same as in the extraction buffer without DTT, PVPP and Sigma protease inhibitor). Total yeast proteins were extracted as described (Kushinov 2000). For immunoblotting analysis, proteins were separated by SDS-PAGE using precast Bis-Tris NuPAGE gels (Life Technologies) and transferred to a nitrocellulose membrane (iBlot 2 transfer stacks, Life Technologies). For immunoblotting assays involving full length RGA5 or RGA5_ΔHMA, proteins were wet-transferred to a nitrocellulose membrane (Millipore). Membranes were blocked in 5% skimmed milk and probed with anti-HA (Roche anti-HA-HRP 3F10), anti-Myc mouse monoclonal antibodies (Roche) or anti-GFP mouse antibodies (Roche) followed, if required, by goat anti-mouse antibodies conjugated with horseradish peroxidase (Sigma). Labeling was detected using the Immobilon western kit (Millipore) or the SuperSignal West Femto Maximum sensitivity substrate (Thermo Fisher Scientific). Membranes were stained with Ponceau S to confirm equal loading.

### • Statistical analysis for *M. oryzae* inoculation test

For each tested *M. oryzae* strain, lesion surfaces were measured on the youngest fully expanded leaf of 5-8 plants per independent transgenic rice line using LeAFtool (https://github.com/sravel/LeAFtool). To determine whether lesion areas on different transgenic lines of rice are significantly different, a Kruskal-Wallis test was performed followed by a Dunn test.

### • Molecular modelling of HMA/AVR complexes

The 3D model of the RGA5_HMA/AVR-Pia complex was generated by replacing the AVR1-CO39 molecule in the RGA5_HMA/AVR1_CO39 crystal structure (PDB 5ZNG) with the superimposed AVR-Pia molecule from the Pikp-1_HMA/AVR-Pia crystal structure (PDB 6Q76). The RGA5_HMAm1m2/AVR1-CO39 3D model was built by replacing in PDB 5ZNG the peptide fragments containing strands β2-β3 (Gly1024-Gly1042) and β4 (Ala1061-Val1069) by the corresponding superimposed peptide fragments from the Pikp-1 HMA molecule in PDB 6G10 (Gly215-Gly233 and Ala252-Lys262, respectively) and substituting Ala260 and Asn261 with Val and Glu in order to match the RGA5_HMAm1m2 C-terminal sequence. The AVR1-CO39 molecule was then replaced by the superimposed AVR-PikD effector molecule in PDB 6G10 in order to model the RGA5_HMAm1m2/AVR-PikD complex. The RGA5_HMAm1m2/AVR-Pia complex was modelled by replacing the effector and peptide fragments from PDB 6G10 in the RGA5_HMAm1m2/AVR-PikD model by their structural counterparts in PDB 6Q76. All models were then refined in explicit water with Charmm in Charmm36 force field (Jo *et al*., 2008). The refinement protocol consisted of 100 energy minimization steps followed by 125 ps (125000 steps of 1fsec) molecular dynamics at 303K, sufficient to reach stable equilibrium as observed by reporting room mean square fluctuations of energies and temperature. Analysis of the HMA/AVR complex interface was performed with QtPISA (Krissinel, 2015).

## ACKNOWLEDGMENTS

We thank Mark Banfield and Hannah Langlands for providing the *AVR-PikD:HA* and *Pikp-1:Flag/Pikp-2:HA* constructs. We thank Corinne Michel and Aurélie Ducasse for technical assistance. This research was funded by the ANR project Immunereceptor (ANR-15-CE20-0007) and benefited from the PhD fellowship of Y.X. from Chinese Scholarship Council (CSC grant 201806350131) and from interactions promoted by COST Action Sustain FA1208 (https://www.cost-sustain.org). The CBS is a member of the French Infrastructure for Integrated Structural Biology (FRISBI), supported by the National Research Agency (<u>ANR-10-INBS-05</u>) and is a GIS-IBIsA platform.

## AUTHOR CONTRIBUTIONS

S.C., A.P. and T.K. designed research. S.C., Y.X., N.D., V.C., L.M., M.P., C.H., K.d.G. and A.P. performed research. S.C., Y.X., N.D., K.d.G., A.P. and T.K. analyzed data. S.C., N.D., and T.K. wrote the paper.

## DATA AVAILABILITY STATEMENT

The datasets generated during and/or analyzed during the current study are available from the corresponding authors on reasonable request.

## SUPPLEMENTAL FIGURE LEGENDS

**Supplemental figure 1: 3D models of engineered RGA5_HMAm1m2 in complex with MAX effectors. A)** Sequence alignment of RGA5 and Pikp-1 HMA domains and of the engineered RGA5_HMAm1m2. The residues constituting the binding interface in the complexes AVR1-CO39/RGA5_HMA (PDB_5ZNG), AVR-Pia/Pikp-1_HMA (PDB_6Q76) and AVR-PikD/Pikp-1_HMA (PDB_6G10) are highlighted respectively in pink, blue and green. The peptide fragments comprising strands β2/β3 and β4, targeted respectively by the m1 and m2 mutations and extracted from Pikp-1_HMA crystal structures to construct chimeric 3D models of RGA5_HMAm1m2, are highlighted in green. **B)** Structural models of RGA5_HMAm1m2 in complex with AVR-PikD (orange), AVR1-CO39 (blue) and AVR-Pia (cyan) are shown as cartoons (top views) or molecular surfaces (orthogonal bottom views). The different MAX effector/RGA5_HMAm1m2 complex structures were built by replacing in PDB_5ZNG the effector molecule and the peptide fragments containing the m1 and m2 mutations by their structural counterpart in PDB_6G10 or PDB_6Q76. The secondary structure elements of RGA5_HMAm1m2 in contact with the MAX effectors are colored as in panel (A). Side chains are shown as sticks for the mutated residues in RGA5_HMAm1m2, and for the residues replaced in the inactive effector variants AVR1-CO39_T41G and AVR-Pia_F24S (both highlighted in red in the bottom views) used as negative control in SPR experiments.

**Supplemental figure 2: The m1m2 mutation does not abolish AVR1-CO39 binding to RGA5_C-ter.** Interaction of BD-fused AVR1-CO39 (without signal peptides) with the AD-fused C-terminal domains of RGA5 and RGA5m1m2 (residues 883 to 1116) was assayed by yeast two-hybrid experiments. The HMA domain of Pikp-1 (AD:Pikp-1_HMA) and the AD and BD domains of GAL4 were used as controls. Four dilutions of diploid yeast clones (1/1, 1/10, 1/100, 1/1000) were spotted on synthetic TDO (-Trp/-Leu/-His) medium and TDO supplemented with 1 mM of 3-amino-1,2,4-triazole (3AT) to assay for interactions and on synthetic DDO (-Trp/-Leu) to monitor proper growth. Pictures were taken after 5 days of growth.

**Supplemental figure 3: Presence and integrity of chimeric proteins expressed in yeast.** Total proteins were extracted from yeasts and Myc- and HA-tagged proteins were detected by immunoblotting using anti-Myc and anti-HA antibodies, respectively.

**Supplemental figure 4. Single cycle kinetic titrations with AVR-PikD to different MBP:HMAs.** The SPR sensorgrams (black curves) show the interaction of the AVR-PikD effector with the different wild-type and mutant HMA domains fused to MBP and captured by anti-MBP antibody immobilized on the chip. Black arrows indicate successive injections of AVR-PikD for 60 sec at the indicated protein concentration, followed by a dissociation phase in running buffer of 600 sec after the final injection. The red curves are the fitting curves obtained with the BiaEvaluation program using a steady-state model (panel A) or a heterogeneous kinetic model (panel B-D). Kinetic data are reported in Supplemental Table 2.

**Supplemental figure 5: Presence and integrity of proteins expressed in *N. benthamiana*.** Immunoblotting showing expression of HA- and YFP-fused proteins. Total proteins were extracted from transiently transformed *N. benthamiana* leaves 48 h after infiltration and were analyzed by immunoblotting with anti-GFP or anti-HA antibodies. Ponceau staining was used to verify equal protein loading.

**Supplemental figure 6: AVR recognition specificities in *N. benthamiana*.** The indicated combinations of constructs were transiently expressed in *N. benthamiana* leaves. Cell death was visualized 5 days after infiltration. Greyscale pictures were taken using a fluorescence scanner with settings allowing visualization of cell death (white patches).

**Supplemental figure 7: Recognition of AVR-Pia and AVR-PikD by RGA5m1, m2 and m1m2.** The indicated combinations of constructs were transiently expressed in *N. benthamiana* leaves. Greyscale pictures were taken using a fluorescence scanner with settings allowing visualization of the disappearance of red fluorescence due to cell death. Therefore, cell death was quantified by measuring fluorescence levels (arbitrary unit, AU) in the infiltrated areas using ImageJ (Xi *et al*., Under review). Resulting data were plotted. The boxes represent the first quartile, median, and third quartile. A Kruskal Wallis test followed by a Dunn test were performed to assess difference of fluorescence levels among the various conditions. Groups with the same letter (A to D) are not significantly different at level 0.01. For each combination of constructs, all of the measurements are represented as dots with a distinct color (red, green and blue) for each of the three biological replicates. The sample size (n) for each experimental group is given.

**Supplemental figure 8: Inoculation of T0 transgenic plants with *M. oryzae*.** The rice cultivar Nipponbare was co-transformed with a genomic construct for *RGA4* and a genomic construct for *RGA5*, *RGA5m1, RGA5m2* or *RGA5m1m2*. A transgenic line carrying *RGA4* and the *GFP* was also generated as a control. T0 plants of the transgenic lines were spray inoculated with the transgenic strain Guy11-AVR-Pia or the wild-type JP10 (*AVR-PikD+*) isolate. The rice cultivar K60 carrying the *Pikp* resistance was used as a control for AVR-PikD specific recognition while Nipponbare (*pikp-/pia-*) served as negative control. Pictures show representative symptoms at 7 days after inoculation. Individual leaves indicate independent T1 transgenic lines (see Supplemental Table 3). S = susceptible, R = resistant.

**Supplemental figure 9: Disease lesion measurements after inoculation of T2 transgenic plants.** Transgenic *M. oryzae* isolates carrying *AVR-Pia*, *AVR-PikD* or the empty vector (EV), were spray-inoculated on T2 transgenic plants carrying the indicated transgenes (i.e. *RGA4+GFP, RGA4+RGA5, RGA4+RGA5m1, RGA4+RGA5m1m2* or *RGA4+RGA5m2*). For each combination of transgenes, the rice transgenic lines used for inoculation are indicated (see Suppl. Table 3). Leaves from 5 to 8 different plants for each transgenic line were scanned 7 days after inoculation. Areas of disease lesions were measured using LeAFtool (https://github.com/sravel/LeAFtool) and plotted. The boxes represent the first quartile, median, and third quartile. Difference of lesion areas among the transgenic lines was assessed by a Kruskall-Wallis test followed by a Dunn test. For each isolate inoculated, groups with the same letter (A to C or D) are not significantly different at level 0.01.

