## Supplemental Information for "Design of a new effector recognition specificity in a plant NLR immune receptor by molecular engineering of its integrated decoy domain"

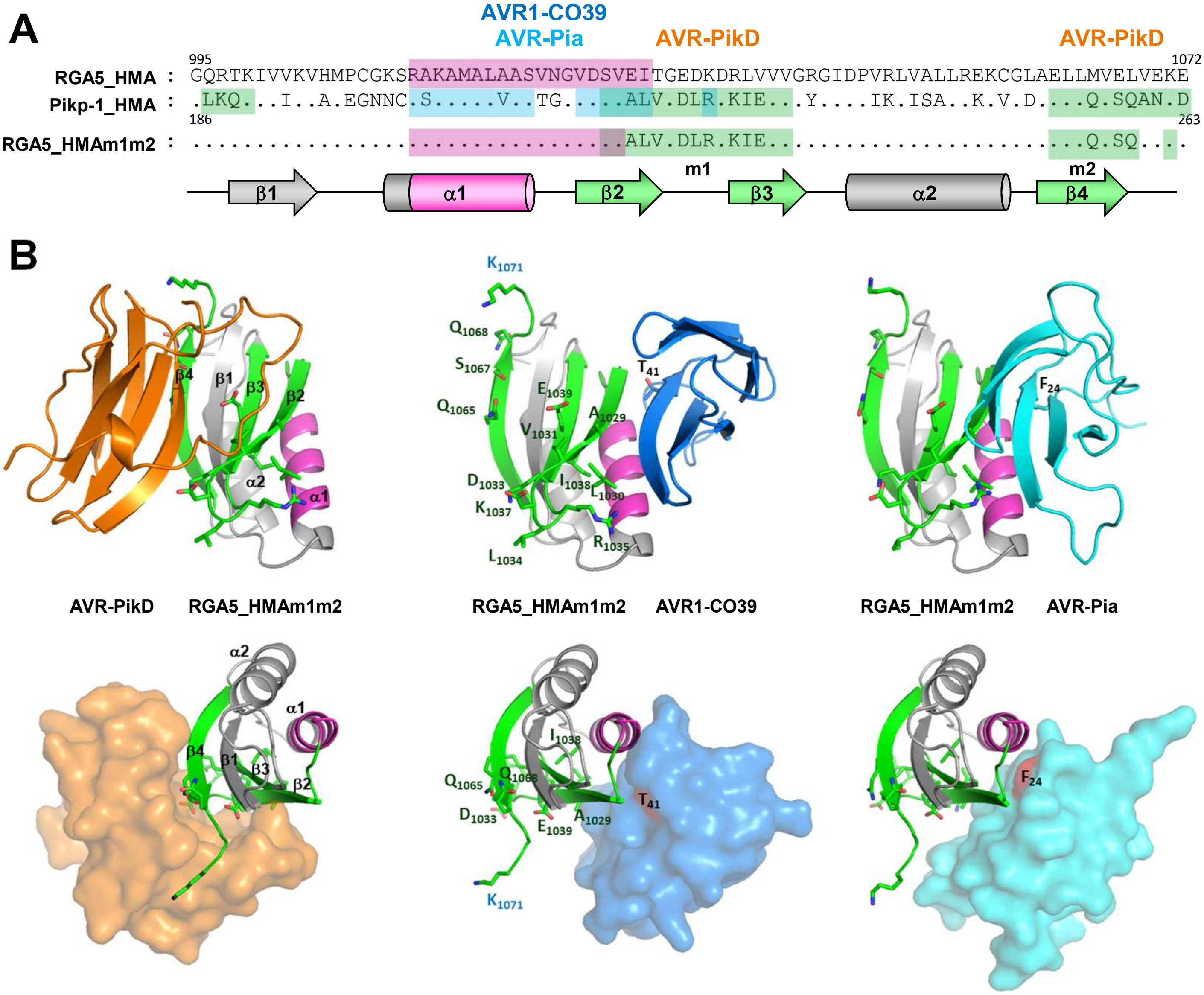


**Supplemental figure 1: 3D models of engineered RGA5_HMAm1m2 in complex with MAX effectors.** **A)** Sequence alignment of RGA5 and Pikp-1 HMA domains and of the engineered RGA5_HMAm1m2. The residues constituting the binding interface in the complexes AVR1-CO39/RGA5_HMA (PDB_5ZNG), AVR-Pia/Pikp-1_HMA (PDB_6Q76) and AVR-PikD/Pikp-1_HMA (PDB_6G10) are highlighted respectively in pink, blue and green. The peptide fragments comprising strands β2/β3 and β4, targeted respectively by the m1 and m2 mutations and extracted from Pikp-1_HMA crystal structures to construct chimeric 3D models of RGA5_HMAm1m2, are highlighted in green. **B)** Structural models of RGA5_HMAm1m2 in complex with AVR-PikD (orange), AVR1-CO39 (blue) and AVR-Pia (cyan) are shown as cartoons (top views) or molecular surfaces (orthogonal bottom views). The different MAX effector/RGA5_HMAm1m2 complex structures were built by replacing in PDB_5ZNG the effector molecule and the peptide fragments containing the m1 and m2 mutations by their structural counterpart in PDB_6G10 or PDB_6Q76. The secondary structure elements of RGA5_HMAm1m2 in contact with the MAX effectors are colored as in panel (A). Side chains are shown as sticks for the mutated residues in RGA5_HMAm1m2, and for the residues replaced in the inactive effector variants AVR1-CO39_T41G and AVR-Pia_F24S (both highlighted in red in the bottom views) used as negative control in SPR experiments.

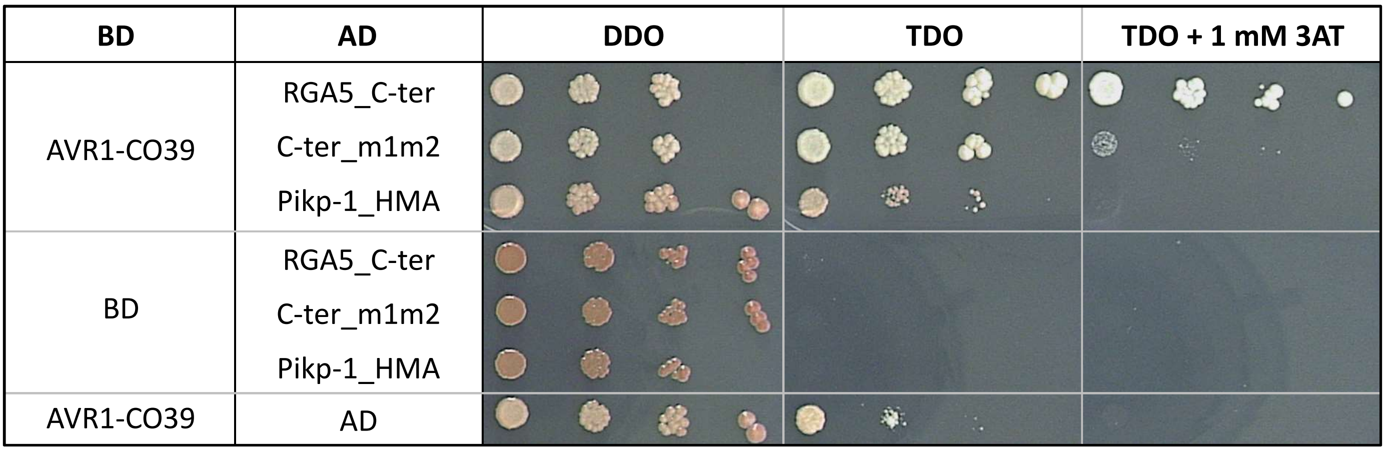


**Supplemental figure 2: The m1m2 mutation does not abolish AVR1-CO39 binding to RGA5_C-ter.** Interaction of BD-fused AVR1-CO39 (without signal peptides) with the AD-fused C-terminal domains of RGA5 and RGA5m1m2 (residues 883 to 1116) was assayed by yeast two-hybrid experiments. The HMA domain of Pikp-1 (AD:Pikp-1_HMA) and the AD and BD domains of GAL4 were used as controls. Four dilutions of diploid yeast clones (1/1, 1/10, 1/100, 1/1000) were spotted on synthetic TDO (-Trp/-Leu/-His) medium and TDO supplemented with 1 mM of 3-amino-1,2,4-triazole (3AT) to assay for interactions and on synthetic DDO (-Trp/-Leu) to monitor proper growth. Pictures were taken after 5 days of growth.

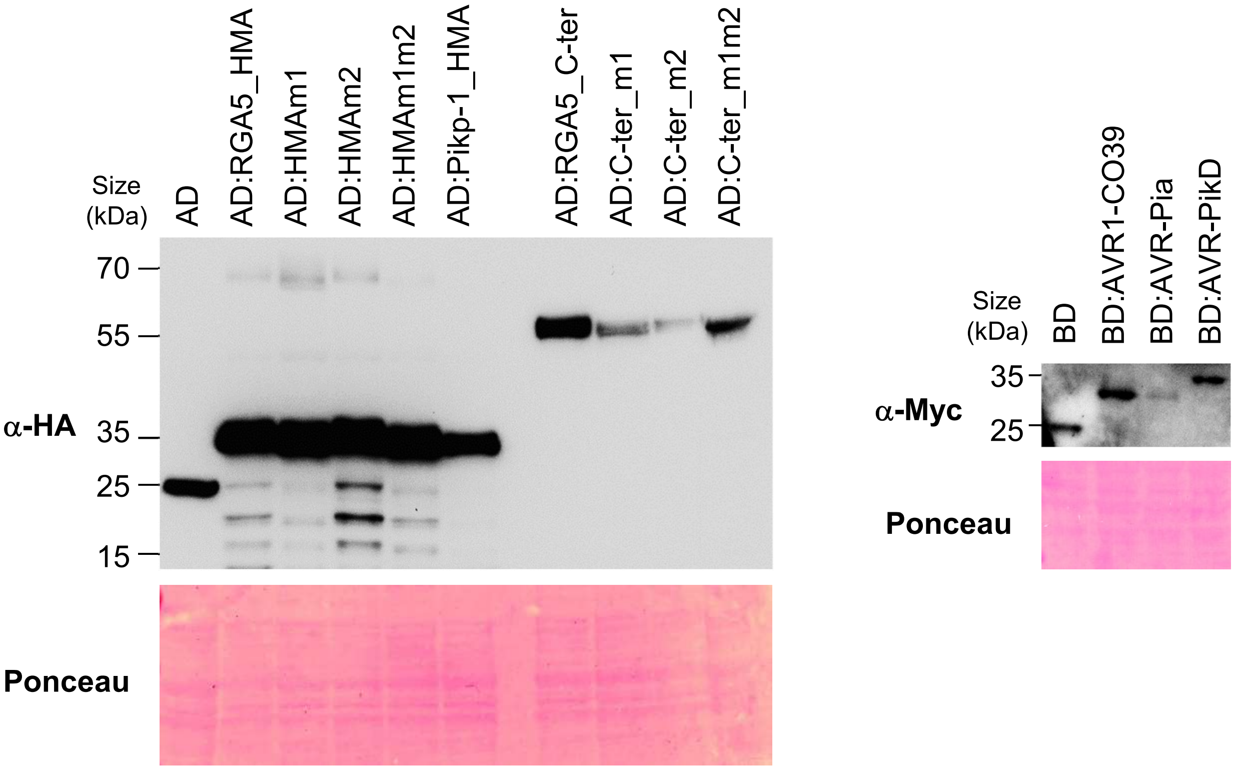


**Supplemental figure 3: Presence and integrity of chimeric proteins expressed in yeast.** Total proteins were extracted from yeasts and Myc- and HA-tagged proteins were detected by immunoblotting using anti-Myc and anti-HA antibodies, respectively.

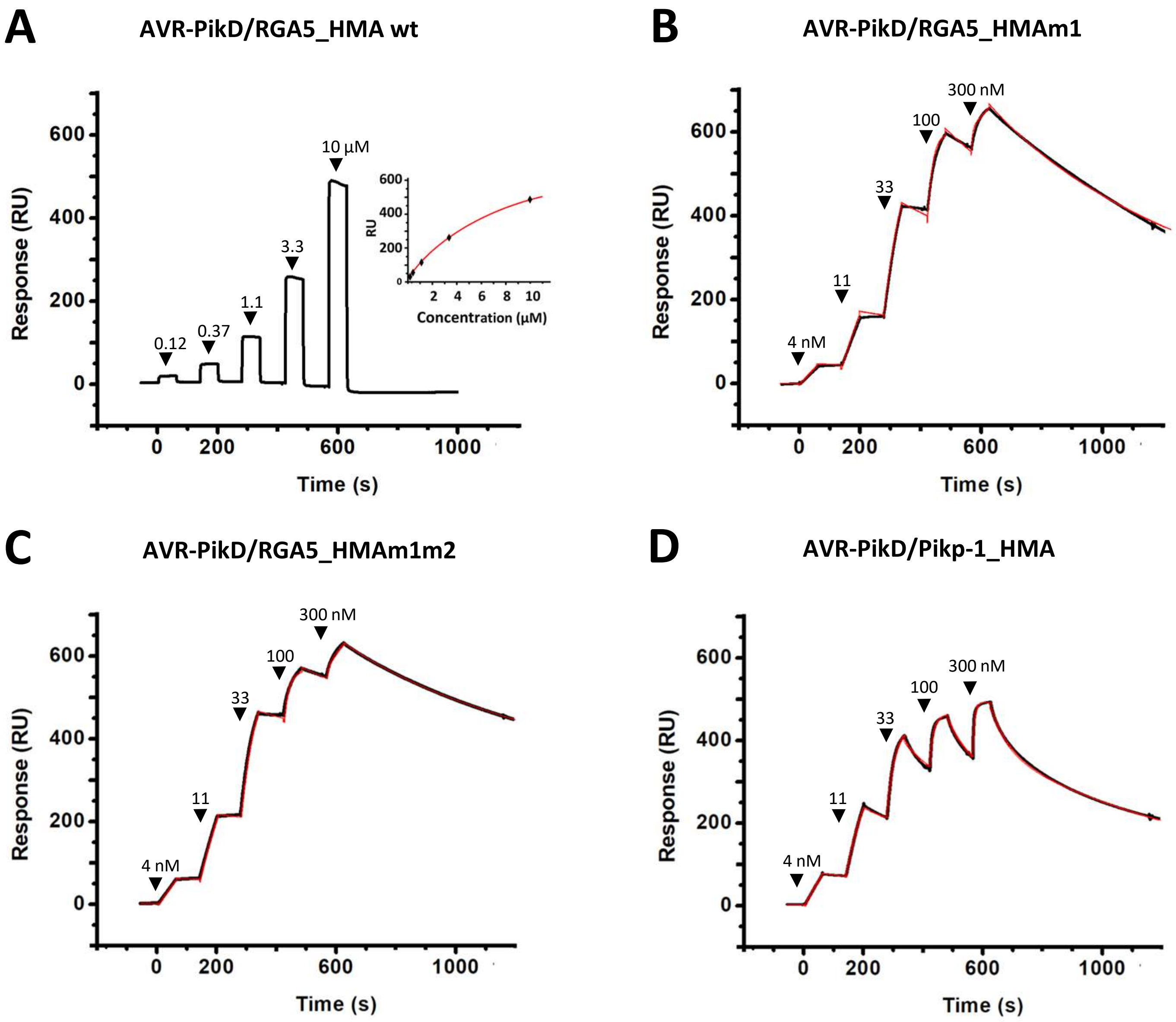


**Supplemental figure 4: Single cycle kinetic titrations with AVR-PikD to different MBP:HMAs.** The SPR sensorgrams (black curves) show the interaction of the AVR-PikD effector with the different wild-type and mutant HMA domains fused to MBP and captured by anti-MBP antibody immobilized on the chip. Black arrows indicate successive injections of AVR-PikD for 60 sec at the indicated protein concentration, followed by a dissociation phase in running buffer of 80 sec or 600 sec for the final injection. The red curves show the data fit performed by the BiaEvaluation program using a steady-state model (panel A) or a heterogeneous kinetic model (panel B-D). Binding and fitting parameters are reported in Supplemental Table 2.

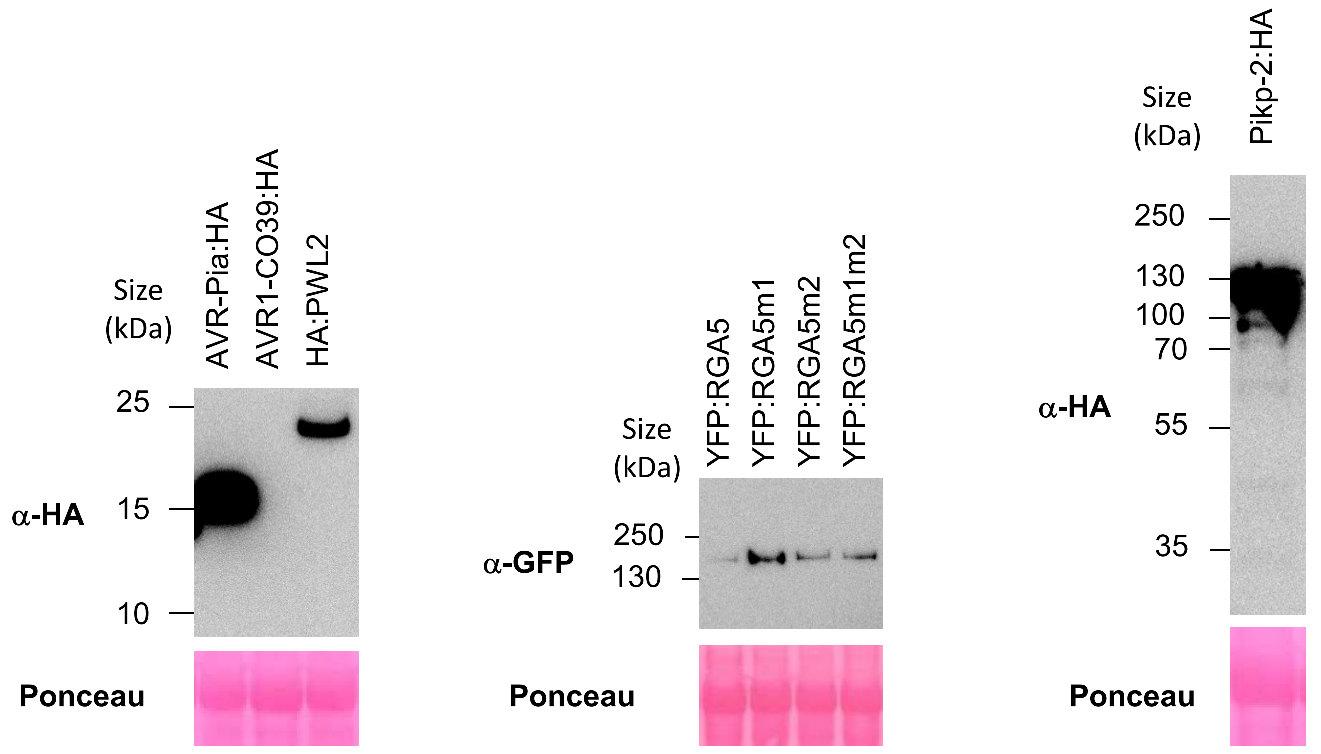


**Supplemental figure 5: Presence and integrity of proteins expressed in *N. benthamiana.*** Immunoblotting showing expression of HA- and YFP-fused proteins. Total proteins were extracted from transiently transformed *N. benthamiana* leaves 48 h after infiltration and were analyzed by immunoblotting with anti-GFP or anti-HA antibodies. Ponceau staining was used to verify equal protein loading.

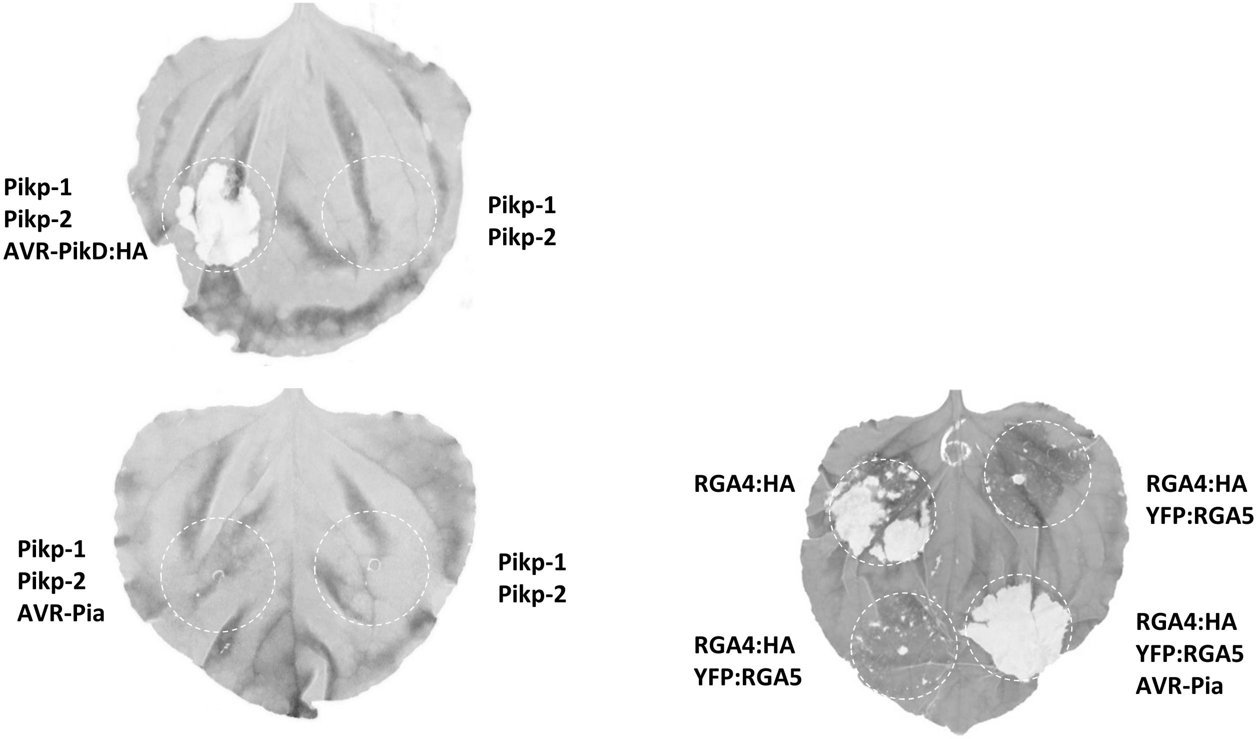


**Supplemental figure 6: AVR recognition specificity in *N. benthamiana*.** The indicated combinations of constructs were transiently expressed in *N. benthamiana* leaves. Cell death was visualized 5 days after infiltration. Greyscale pictures were taken using a fluorescence scanner with settings allowing visualization of cell death (white patches).

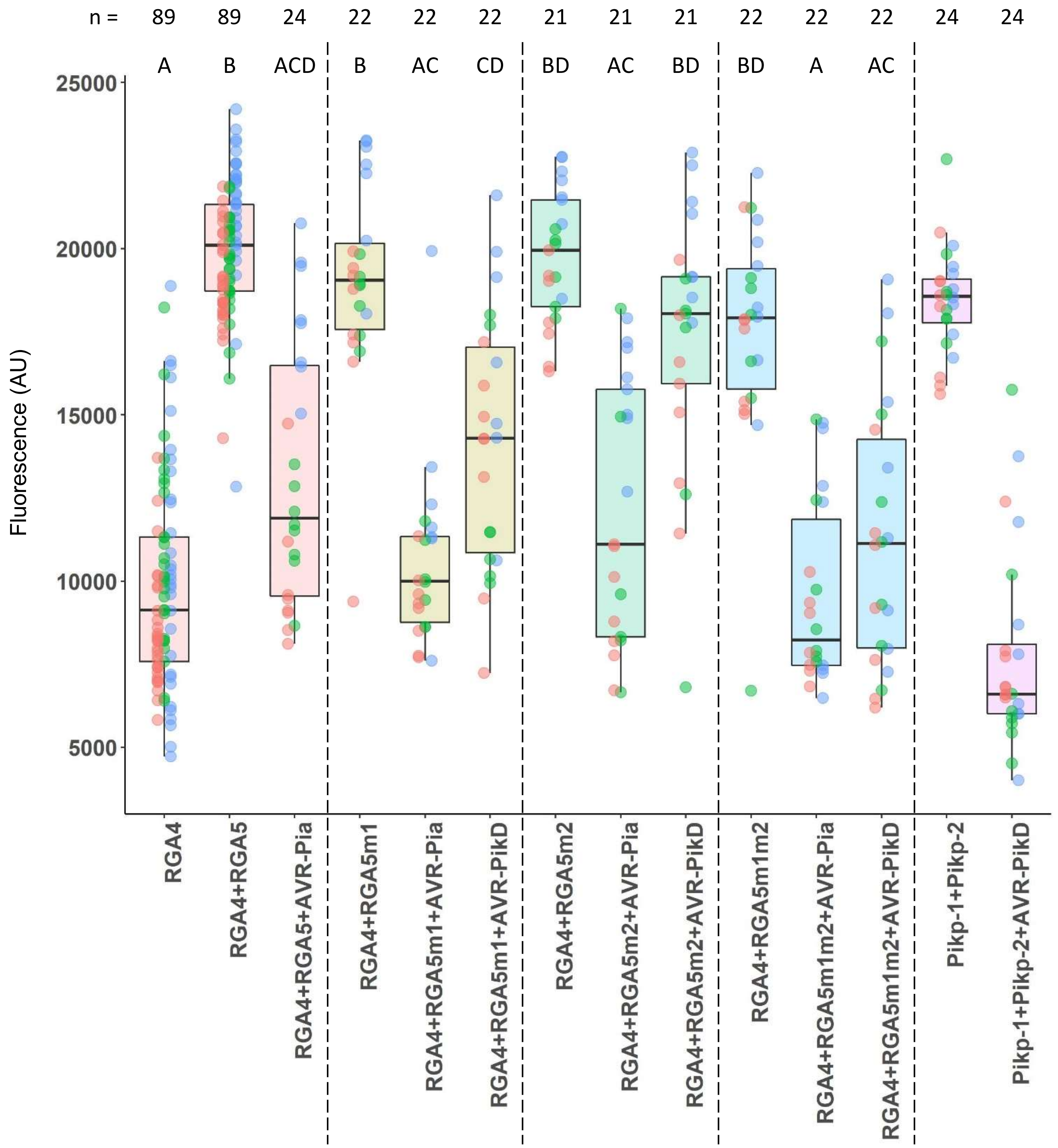


**Supplemental figure 7: Recognition of AVR-Pia and AVR-PikD by RGA5m1, m2 and m1m2.** The indicated combinations of constructs were transiently expressed in *N. benthamiana* leaves. Greyscale pictures were taken using a fluorescence scanner with settings allowing visualization of the disappearance of red fluorescence due to cell death. Therefore, cell death was quantified by measuring fluorescence levels (arbitrary unit, AU) in the infiltrated areas using ImageJ (Xi et al., Under review). Resulting data were plotted. The boxes represent the first quartile, median, and third quartile. A Kruskal Wallis test followed by a Dunn test were performed to assess difference of fluorescence levels among the various conditions. Groups with the same letter (A to D) are not significantly different at level 0.01. For each combination of constructs, all of the measurements are represented as dots with a distinct color (red, green and blue) for each of the three biological replicates. The sample size (n) for each experimental group is given.

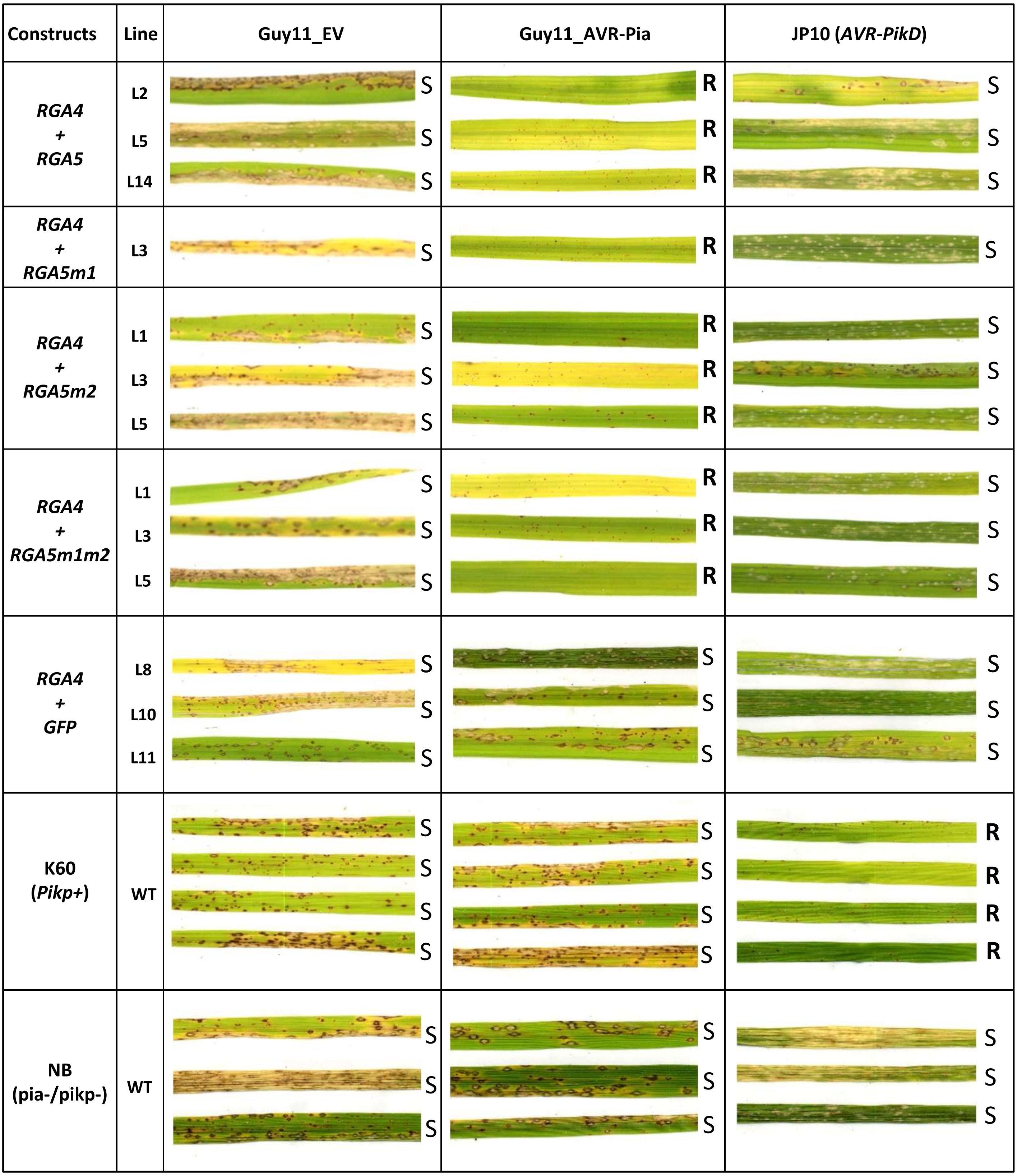


**Supplemental figure 8: Inoculation of T0 transgenic plants with *M. oryzae*.** The rice cultivar Nipponbare was co-transformed with a genomic construct for *RGA4* and a genomic construct for *RGA5*, *RGA5m1*, *RGA5m2* or *RGA5m1m2*. A transgenic line carrying *RGA4* and the *GFP* was also generated as a control. T0 plants of the transgenic lines were spray inoculated with the transgenic strain Guy11-AVR-Pia or the wild-type JP10 (*AVR-PikD*+) isolate. The rice cultivar K60 carrying the *Pikp* resistance was used as a control for AVR-PikD specific recognition while Nipponbare (*pikp-/pia-*) served as negative control. Pictures show representative symptoms at 7 days after inoculation. Individual leaves indicate independent T1 transgenic lines (see Supplemental Table 3). S = susceptible, R = resistant.

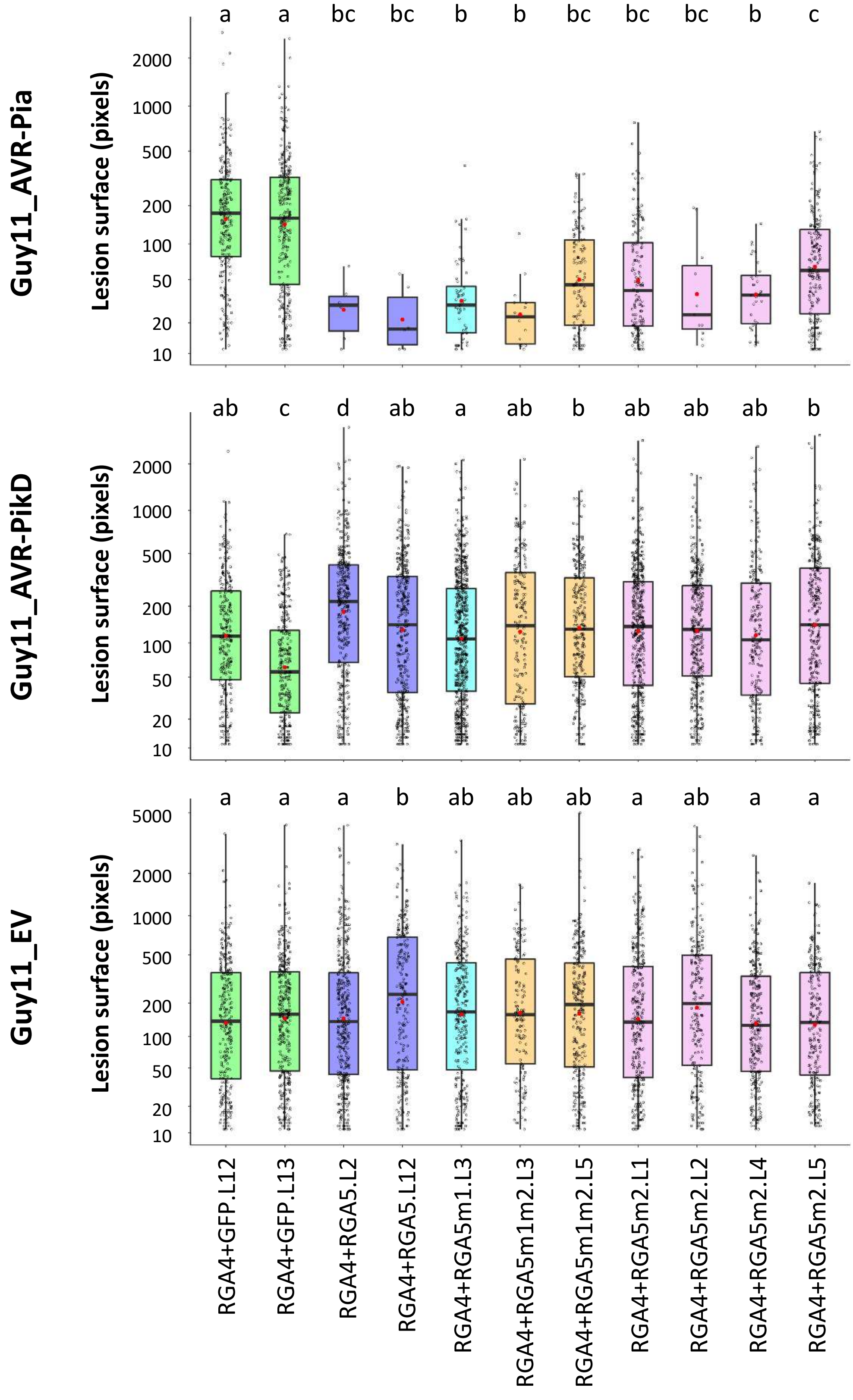


**Supplemental figure 9: Disease lesion measurements after inoculation of T2 transgenic plants.** Transgenic *M. oryzae* isolates carrying *AVR-Pia*, *AVR-PikD* or the empty vector (EV), were spray-inoculated on T2 transgenic plants carrying the indicated transgenes (i.e. *RGA4+GFP*, *RGA4+RGA5, RGA4+RGA5m1, RGA4+RGA5m1m2* or *RGA4+RGA5m2*). For each combination of transgenes, the rice transgenic lines used for inoculation are indicated (see Suppl. Table 3). Leaves from 5 to 8 different plants for each transgenic line were scanned 7 days after inoculation. Areas of disease lesions were measured using LeAFtool (https://github.com/sravel/LeAFtool) and plotted. The boxes represent the first quartile, median, and third quartile. Difference of lesion areas among the transgenic lines was assessed by a Kruskall-Wallis test followed by a Dunn test. For each isolate inoculated, groups with the same letter (A to C or D) are not significantly different at level 0.01.

**Supplemental text 1: Nucleotide sequence of synthetic genes and pDB_ccdb derivatives used in this study**

>*AVR-PikD*
TCCAGGGGCCCCATATGGAGACGGGTAATAAGTACATCGAGAAACGTGCGATCGACCTTTCACGCG
AACGCGACCCGAACTTTTTCGATCATCCGGGCATTCCAGTCCCAGAATGCTTCTGGTTTATGTTTAAG
AATAATGTCCGCCAAGACGCAGGAACGTGCTATTCTTCGTGGAAAATGGACATGAAAGTTGGACCCA
ATTGGGTCCATATTAAATCTGACGATAACTGTAACTTGAGCGGGGACTTCCCTCCCGGATGGATCGT
GCTTGGTAAGAAGCGCCCTGGTTTCCTTGAATAATGACTCGAGCACCACCACC

>*AVR1-CO39_T41G*
TCCAGGGGCCCCATATGGCATGGAAAGATTGCATTATTCAACGTTACAAAGATGGAGACGTTAACAA
CATCTACGGGGCCAATCGCAACGAAGAGATCACCATCGAGGAATATAAGGTTTTCGTCAATGAAGCC
TGCCACCCGTACCCGGTGATCCTGCCCGATAAAAGTGTTTTATCGGGGGACTTCACAAGCGCATACG
CGGACGATGACGAGTCATGCTAATGACTCGAGCACCACCACC

>*RGA5_HMA*
TCCAGGGGCCCCATATGAGTGCATTAACGGGGCAACGAACTAAGATAGTTGTTAAGGTGCACATGC
CATGCGGAAAATCCCGAGCAAAAGCCATGGCGCTGGCTGCGTCAGTGAACGGGGTGGACAGCGTG
GAGATAACGGGGGAGGACAAAGACCGGCTGGTGGTGGTCGGCCGTGGCATTGACCCTGTTCGCCT
GGTGGCTCTCCTGCGCGAGAAATGTGGCCTCGCCGAGCTCTTGATGGTGGAGTTAGTTGAGAAAGA
GTGATAATGACTCGAGCACCACCACC

> *RGA5_HMAm1*
TCCAGGGGCCCCATATGAGTGCATTAACGGGGCAACGAACTAAGATAGTTGTTAAGGTGCACATGC
CATGCGGAAAATCCCGAGCAAAAGCCATGGCGCTGGCTGCGTCAGTGAACGGGGTGGACAGCGTG
GCGTTAGTGGGGGATCTAAGAGACAAGATCGAGGTGGTCGGCCGTGGCATTGACCCTGTTCGCCTG
GTGGCTCTCCTGCGCGAGAAATGTGGCCTCGCCGAGCTCTTGATGGTGGAGTTAGTTGAGAAAGAG
TGATAATGACTCGAGCACCACCACC

> *RGA5_HMAm1m2*
TCCAGGGGCCCCATATGAGTGCATTAACGGGGCAACGAACTAAGATAGTTGTTAAGGTGCACATGC
CATGCGGAAAATCCCGAGCAAAAGCCATGGCGCTGGCTGCGTCAGTGAACGGGGTGGACAGCGTG
GCGTTAGTGGGGGATCTAAGAGACAAGATCGAGGTGGTCGGCCGTGGCATTGACCCTGTTCGCCTG
GTGGCTCTCCTGCGCGAGAAATGTGGCCTCGCCGAGCTCTTGCAGGTGTCGCAGGTTGAGAAAGAG
TGATAATGACTCGAGCACCACCACC

> *Pikp-1_HMA*
TCCAGGGGCCCcatatgATGAGCGATTACGACATCCCCACTACTAAGCTTCTGGAAGTTCTGTTCCAGG
GGCCCCATATGGGCCTGAAGCAGAAGATTGTAATTAAAGTTGCTATGGAAGGCAACAACTGTCGCA
GTAAAGCAATGGCGCTGGTAGCGTCCACGGGCGGGGTTGACTCAGTGGCTCTTGTAGGTGATTTGC
GTGATAAAATCGAGGTGGTAGGGTATGGTATTGACCCTATCAAGCTGATCTCAGCGTTGCGCAAAAA
AGTTGGCGACGCGGAACTGTTGCAGGTCAGCTAATGACTCGAGCACCACCACC

> pDB_ccdb_his_3C
TGGCGAATGGGACGCGCCCTGTAGCGGCGCATTAAGCGCGGCGGGTGTGGTGGTTACGCGCAGCG
TGACCGCTACACTTGCCAGCGCCCTAGCGCCCGCTCCTTTCGCTTTCTTCCCTTCCTTTCTCGCCACGTT

CGCCGGCTTTCCCCGTCAAGCTCTAAATCGGGGGCTCCCTTTAGGGTTCCGATTTAGTGCTTTACGGC
ACCTCGACCCCAAAAAACTTGATTAGGGTGATGGTTCACGTAGTGGGCCATCGCCCTGATAGACGGT
TTTTCGCCCTTTGACGTTGGAGTCCACGTTCTTTAATAGTGGACTCTTGTTCCAAACTGGAACAACACT
CAACCCTATCTCGGTCTATTCTTTTGATTTATAAGGGATTTTGCCGATTTCGGCCTATTGGTTAAAAAA
TGAGCTGATTTAACAAAAATTTAACGCGAATTTTAACAAAATATTAACGTTTACAATTTCAGGTGGCA
CTTTTCGGGGAAATGTGCGCGGAACCCCTATTTGTTTATTTTTCTAAATACATTCAAATATGTATCCGC
TCATGAATTAATTCTTAGAAAAACTCATCGAGCATCAAATGAAACTGCAATTTATTCATATCAGGATT
ATCAATACCATATTTTTGAAAAAGCCGTTTCTGTAATGAAGGAGAAAACTCACCGAGGCAGTTCCATA
GGATGGCAAGATCCTGGTATCGGTCTGCGATTCCGACTCGTCCAACATCAATACAACCTATTAATTTC
CCCTCGTCAAAAATAAGGTTATCAAGTGAGAAATCACCATGAGTGACGACTGAATCCGGTGAGAATG
GCAAAAGTTTATGCATTTCTTTCCAGACTTGTTCAACAGGCCAGCCATTACGCTCGTCATCAAAATCAC
TCGCATCAACCAAACCGTTATTCATTCGTGATTGCGCCTGAGCGAGACGAAATACGCGATCGCTGTTA
AAAGGACAATTACAAACAGGAATCGAATGCAACCGGCGCAGGAACACTGCCAGCGCATCAACAATA
TTTTCACCTGAATCAGGATATTCTTCTAATACCTGGAATGCTGTTTTCCCGGGGATCGCAGTGGTGAG
TAACCATGCATCATCAGGAGTACGGATAAAATGCTTGATGGTCGGAAGAGGCATAAATTCCGTCAGC
CAGTTTAGTCTGACCATCTCATCTGTAACATCATTGGCAACGCTACCTTTGCCATGTTTCAGAAACAAC
TCTGGCGCATCGGGCTTCCCATACAATCGATAGATTGTCGCACCTGATTGCCCGACATTATCGCGAGC
CCATTTATACCCATATAAATCAGCATCCATGTTGGAATTTAATCGCGGCCTAGAGCAAGACGTTTCCC
GTTGAATATGGCTCATAACACCCCTTGTATTACTGTTTATGTAAGCAGACAGTTTTATTGTTCATGACC
AAAATCCCTTAACGTGAGTTTTCGTTCCACTGAGCGTCAGACCCCGTAGAAAAGATCAAAGGATCTTC
TTGAGATCCTTTTTTTCTGCGCGTAATCTGCTGCTTGCAAACAAAAAAACCACCGCTACCAGCGGTGG
TTTGTTTGCCGGATCAAGAGCTACCAACTCTTTTTCCGAAGGTAACTGGCTTCAGCAGAGCGCAGATA
CCAAATACTGTCCTTCTAGTGTAGCCGTAGTTAGGCCACCACTTCAAGAACTCTGTAGCACCGCCTAC
ATACCTCGCTCTGCTAATCCTGTTACCAGTGGCTGCTGCCAGTGGCGATAAGTCGTGTCTTACCGGGT
TGGACTCAAGACGATAGTTACCGGATAAGGCGCAGCGGTCGGGCTGAACGGGGGGTTCGTGCACA
CAGCCCAGCTTGGAGCGAACGACCTACACCGAACTGAGATACCTACAGCGTGAGCTATGAGAAAGC
GCCACGCTTCCCGAAGGGAGAAAGGCGGACAGGTATCCGGTAAGCGGCAGGGTCGGAACAGGAGA
GCGCACGAGGGAGCTTCCAGGGGGAAACGCCTGGTATCTTTATAGTCCTGTCGGGTTTCGCCACCTC
TGACTTGAGCGTCGATTTTTGTGATGCTCGTCAGGGGGGCGGAGCCTATGGAAAAACGCCAGCAAC
GCGGCCTTTTTACGGTTCCTGGCCTTTTGCTGGCCTTTTGCTCACATGTTCTTTCCTGCGTTATCCCCTG
ATTCTGTGGATAACCGTATTACCGCCTTTGAGTGAGCTGATACCGCTCGCCGCAGCCGAACGACCGA
GCGCAGCGAGTCAGTGAGCGAGGAAGCGGAAGAGCGCCTGATGCGGTATTTTCTCCTTACGCATCT
GTGCGGTATTTCACACCGCATATATGGTGCACTCTCAGTACAATCTGCTCTGATGCCGCATAGTTAAG
CCAGTATACACTCCGCTATCGCTACGTGACTGGGTCATGGCTGCGCCCCGACACCCGCCAACACCCG
CTGACGCGCCCTGACGGGCTTGTCTGCTCCCGGCATCCGCTTACAGACAAGCTGTGACCGTCTCCGG
GAGCTGCATGTGTCAGAGGTTTTCACCGTCATCACCGAAACGCGCGAGGCAGCTGCGGTAAAGCTC
ATCAGCGTGGTCGTGAAGCGATTCACAGATGTCTGCCTGTTCATCCGCGTCCAGCTCGTTGAGTTTCT
CCAGAAGCGTTAATGTCTGGCTTCTGATAAAGCGGGCCATGTTAAGGGCGGTTTTTTCCTGTTTGGTC
ACTGATGCCTCCGTGTAAGGGGGATTTCTGTTCATGGGGGTAATGATACCGATGAAACGAGAGAGG
ATGCTCACGATACGGGTTACTGATGATGAACATGCCCGGTTACTGGAACGTTGTGAGGGTAAACAAC
TGGCGGTATGGATGCGGCGGGACCAGAGAAAAATCACTCAGGGTCAATGCCAGCGCTTCGTTAATA
CAGATGTAGGTGTTCCACAGGGTAGCCAGCAGCATCCTGCGATGCAGATCCGGAACATAATGGTGC
AGGGCGCTGACTTCCGCGTTTCCAGACTTTACGAAACACGGAAACCGAAGACCATTCATGTTGTTGC
TCAGGTCGCAGACGTTTTGCAGCAGCAGTCGCTTCACGTTCGCTCGCGTATCGGTGATTCATTCTGCT
AACCAGTAAGGCAACCCCGCCAGCCTAGCCGGGTCCTCAACGACAGGAGCACGATCATGCGCACCC
GTGGGGCCGCCATGCCGGCGATAATGGCCTGCTTCTCGCCGAAACGTTTGGTGGCGGGACCAGTGA
CGAAGGCTTGAGCGAGGGCGTGCAAGATTCCGAATACCGCAAGCGACAGGCCGATCATCGTCGCGC

TCCAGCGAAAGCGGTCCTCGCCGAAAATGACCCAGAGCGCTGCCGGCACCTGTCCTACGAGTTGCAT
GATAAAGAAGACAGTCATAAGTGCGGCGACGATAGTCATGCCCCGCGCCCACCGGAAGGAGCTGAC
TGGGTTGAAGGCTCTCAAGGGCATCGGTCGAGATCCCGGTGCCTAATGAGTGAGCTAACTTACATTA
ATTGCGTTGCGCTCACTGCCCGCTTTCCAGTCGGGAAACCTGTCGTGCCAGCTGCATTAATGAATCGG
CCAACGCGCGGGGAGAGGCGGTTTGCGTATTGGGCGCCAGGGTGGTTTTTCTTTTCACCAGTGAGA
CGGGCAACAGCTGATTGCCCTTCACCGCCTGGCCCTGAGAGAGTTGCAGCAAGCGGTCCACGCTGGT
TTGCCCCAGCAGGCGAAAATCCTGTTTGATGGTGGTTAACGGCGGGATATAACATGAGCTGTCTTCG
GTATCGTCGTATCCCACTACCGAGATATCCGCACCAACGCGCAGCCCGGACTCGGTAATGGCGCGCA
TTGCGCCCAGCGCCATCTGATCGTTGGCAACCAGCATCGCAGTGGGAACGATGCCCTCATTCAGCAT
TTGCATGGTTTGTTGAAAACCGGACATGGCACTCCAGTCGCCTTCCCGTTCCGCTATCGGCTGAATTT
GATTGCGAGTGAGATATTTATGCCAGCCAGCCAGACGCAGACGCGCCGAGACAGAACTTAATGGGC
CCGCTAACAGCGCGATTTGCTGGTGACCCAATGCGACCAGATGCTCCACGCCCAGTCGCGTACCGTC
TTCATGGGAGAAAATAATACTGTTGATGGGTGTCTGGTCAGAGACATCAAGAAATAACGCCGGAAC
ATTAGTGCAGGCAGCTTCCACAGCAATGGCATCCTGGTCATCCAGCGGATAGTTAATGATCAGCCCA
CTGACGCGTTGCGCGAGAAGATTGTGCACCGCCGCTTTACAGGCTTCGACGCCGCTTCGTTCTACCAT
CGACACCACCACGCTGGCACCCAGTTGATCGGCGCGAGATTTAATCGCCGCGACAATTTGCGACGGC
GCGTGCAGGGCCAGACTGGAGGTGGCAACGCCAATCAGCAACGACTGTTTGCCCGCCAGTTGTTGT
GCCACGCGGTTGGGAATGTAATTCAGCTCCGCCATCGCCGCTTCCACTTTTTCCCGCGTTTTCGCAGA
AACGTGGCTGGCCTGGTTCACCACGCGGGAAACGGTCTGATAAGAGACACCGGCATACTCTGCGAC
ATCGTATAACGTTACTGGTTTCACATTCACCACCCTGAATTGACTCTCTTCCGGGCGCTATCATGCCAT
ACCGCGAAAGGTTTTGCGCCATTCGATGGTGTCCGGGATCTCGACGCTCTCCCTTATGCGACTCCTGC
ATTAGGAAGCAGCCCAGTAGTAGGTTGAGGCCGTTGAGCACCGCCGCCGCAAGGAATGGTGCATGC
AAGGAGATGGCGCCCAACAGTCCCCCGGCCACGGGGCCTGCCACCATACCCACGCCGAAACAAGCG
CTCATGAGCCCGAAGTGGCGAGCCCGATCTTCCCCATCGGTGATGTCGGCGATATAGGCGCCAGCAA
CCGCACCTGTGGCGCCGGTGATGCCGGCCACGATGCGTCCGGCGTAGAGGATCGAGATCTCGATCC
CGCGAAATTAATACGACTCACTATAGGGGAATTGTGAGCGGATAACAATTCCCCTCTAGAAATAATT
TTGATTTAACTTTAAGAAGGAGATATACCATGAAACATCACCATCACCATCACCCCATGAGCGATTAC
GACATCCCCACTACTAAGCTTCTGGAAGTTCTGTTCCAGGGGCCCCATATGACTGGCTGTGTATAAGG
GAGCCTGACATTTATATTCCCCAGAACATCAGGTTAATGGCGTTTTTGATGTCATTTTCGCGGTGGCT
GAGATCAGCCACTTCTTCCCCGATAACGGAGACCGGCACACTGGCCATATCGGTGGTCATCATGCGC
CAGCTTTCATCCCCGATATGCACCACCGGGTAAAGTTCACGGGAGACTTTATCTGACAGCAGACGTG
CACTGGCCAGGGGGATCACCATCCGTCGCCCGGGCGTGTCAATAATATCACTCTGTACATCCACAAA
CAGACGATAACGGCTCTCTCTTTTATAGGTGTAAACCTTAAACTGCATTTCACCAGCCCCTGTTCTCGT
CAGCAAAAGAGCCGTTCATTTCAATAAACCGGGCGACCTCAGCCATCCCTTCCTGATTTTCCGCTTTCC
AGCGTTCGGCACGCAGACGACGGGCTTCATTCTGCATGGTTGTGCTTACCAGACCGGAGATATTGAC
ATCATATATGCCTTGAGCAACTGATAGCTGTCGCTGTCAACTGTCACTGTAATACGCTGCTTCATAGC
ATACCTCTTTTTGACATACTTCGGGTATACATATCAGTATATATTCTTATACCGCAAAAATCAGCGCGC
AAATACGCATACTGTTATCTGGCTTTTAGTAAGCCGGATCCACGCGTCTCGAGCACCACCACCACCAC
CACTGAGATCCGGCTGCTAACAAAGCCCGAAAGGAAGCTGAGTTGGCTGCTGCCACCGCTGAGCAA
TAACTAGCATAACCCCTTGGGGCCTCTAAACGGGTCTTGAGGGGTTTTTTGCTGAAAGGAGGAACTA
TATCCGGAT

> pDb_ccdb_pepL_his_3C
TGGCGAATGGGACGCGCCCTGTAGCGGCGCATTAAGCGCGGCGGGTGTGGTGGTTACGCGCAGCG
TGACCGCTACACTTGCCAGCGCCCTAGCGCCCGCTCCTTTCGCTTTCTTCCCTTCCTTTCTCGCCACGTT
CGCCGGCTTTCCCCGTCAAGCTCTAAATCGGGGGCTCCCTTTAGGGTTCCGATTTAGTGCTTTACGGC
ACCTCGACCCCAAAAAACTTGATTAGGGTGATGGTTCACGTAGTGGGCCATCGCCCTGATAGACGGT

TTTTCGCCCTTTGACGTTGGAGTCCACGTTCTTTAATAGTGGACTCTTGTTCCAAACTGGAACAACACT
CAACCCTATCTCGGTCTATTCTTTTGATTTATAAGGGATTTTGCCGATTTCGGCCTATTGGTTAAAAAA
TGAGCTGATTTAACAAAAATTTAACGCGAATTTTAACAAAATATTAACGTTTACAATTTCAGGTGGCA
CTTTTCGGGGAAATGTGCGCGGAACCCCTATTTGTTTATTTTTCTAAATACATTCAAATATGTATCCGC
TCATGAATTAATTCTTAGAAAAACTCATCGAGCATCAAATGAAACTGCAATTTATTCATATCAGGATT
ATCAATACCATATTTTTGAAAAAGCCGTTTCTGTAATGAAGGAGAAAACTCACCGAGGCAGTTCCATA
GGATGGCAAGATCCTGGTATCGGTCTGCGATTCCGACTCGTCCAACATCAATACAACCTATTAATTTC
CCCTCGTCAAAAATAAGGTTATCAAGTGAGAAATCACCATGAGTGACGACTGAATCCGGTGAGAATG
GCAAAAGTTTATGCATTTCTTTCCAGACTTGTTCAACAGGCCAGCCATTACGCTCGTCATCAAAATCAC
TCGCATCAACCAAACCGTTATTCATTCGTGATTGCGCCTGAGCGAGACGAAATACGCGATCGCTGTTA
AAAGGACAATTACAAACAGGAATCGAATGCAACCGGCGCAGGAACACTGCCAGCGCATCAACAATA
TTTTCACCTGAATCAGGATATTCTTCTAATACCTGGAATGCTGTTTTCCCGGGGATCGCAGTGGTGAG
TAACCATGCATCATCAGGAGTACGGATAAAATGCTTGATGGTCGGAAGAGGCATAAATTCCGTCAGC
CAGTTTAGTCTGACCATCTCATCTGTAACATCATTGGCAACGCTACCTTTGCCATGTTTCAGAAACAAC
TCTGGCGCATCGGGCTTCCCATACAATCGATAGATTGTCGCACCTGATTGCCCGACATTATCGCGAGC
CCATTTATACCCATATAAATCAGCATCCATGTTGGAATTTAATCGCGGCCTAGAGCAAGACGTTTCCC
GTTGAATATGGCTCATAACACCCCTTGTATTACTGTTTATGTAAGCAGACAGTTTTATTGTTCATGACC
AAAATCCCTTAACGTGAGTTTTCGTTCCACTGAGCGTCAGACCCCGTAGAAAAGATCAAAGGATCTTC
TTGAGATCCTTTTTTTCTGCGCGTAATCTGCTGCTTGCAAACAAAAAAACCACCGCTACCAGCGGTGG
TTTGTTTGCCGGATCAAGAGCTACCAACTCTTTTTCCGAAGGTAACTGGCTTCAGCAGAGCGCAGATA
CCAAATACTGTCCTTCTAGTGTAGCCGTAGTTAGGCCACCACTTCAAGAACTCTGTAGCACCGCCTAC
ATACCTCGCTCTGCTAATCCTGTTACCAGTGGCTGCTGCCAGTGGCGATAAGTCGTGTCTTACCGGGT
TGGACTCAAGACGATAGTTACCGGATAAGGCGCAGCGGTCGGGCTGAACGGGGGGTTCGTGCACA
CAGCCCAGCTTGGAGCGAACGACCTACACCGAACTGAGATACCTACAGCGTGAGCTATGAGAAAGC
GCCACGCTTCCCGAAGGGAGAAAGGCGGACAGGTATCCGGTAAGCGGCAGGGTCGGAACAGGAGA
GCGCACGAGGGAGCTTCCAGGGGGAAACGCCTGGTATCTTTATAGTCCTGTCGGGTTTCGCCACCTC
TGACTTGAGCGTCGATTTTTGTGATGCTCGTCAGGGGGGCGGAGCCTATGGAAAAACGCCAGCAAC
GCGGCCTTTTTACGGTTCCTGGCCTTTTGCTGGCCTTTTGCTCACATGTTCTTTCCTGCGTTATCCCCTG
ATTCTGTGGATAACCGTATTACCGCCTTTGAGTGAGCTGATACCGCTCGCCGCAGCCGAACGACCGA
GCGCAGCGAGTCAGTGAGCGAGGAAGCGGAAGAGCGCCTGATGCGGTATTTTCTCCTTACGCATCT
GTGCGGTATTTCACACCGCATATATGGTGCACTCTCAGTACAATCTGCTCTGATGCCGCATAGTTAAG
CCAGTATACACTCCGCTATCGCTACGTGACTGGGTCATGGCTGCGCCCCGACACCCGCCAACACCCG
CTGACGCGCCCTGACGGGCTTGTCTGCTCCCGGCATCCGCTTACAGACAAGCTGTGACCGTCTCCGG
GAGCTGCATGTGTCAGAGGTTTTCACCGTCATCACCGAAACGCGCGAGGCAGCTGCGGTAAAGCTC
ATCAGCGTGGTCGTGAAGCGATTCACAGATGTCTGCCTGTTCATCCGCGTCCAGCTCGTTGAGTTTCT
CCAGAAGCGTTAATGTCTGGCTTCTGATAAAGCGGGCCATGTTAAGGGCGGTTTTTTCCTGTTTGGTC
ACTGATGCCTCCGTGTAAGGGGGATTTCTGTTCATGGGGGTAATGATACCGATGAAACGAGAGAGG
ATGCTCACGATACGGGTTACTGATGATGAACATGCCCGGTTACTGGAACGTTGTGAGGGTAAACAAC
TGGCGGTATGGATGCGGCGGGACCAGAGAAAAATCACTCAGGGTCAATGCCAGCGCTTCGTTAATA
CAGATGTAGGTGTTCCACAGGGTAGCCAGCAGCATCCTGCGATGCAGATCCGGAACATAATGGTGC
AGGGCGCTGACTTCCGCGTTTCCAGACTTTACGAAACACGGAAACCGAAGACCATTCATGTTGTTGC
TCAGGTCGCAGACGTTTTGCAGCAGCAGTCGCTTCACGTTCGCTCGCGTATCGGTGATTCATTCTGCT
AACCAGTAAGGCAACCCCGCCAGCCTAGCCGGGTCCTCAACGACAGGAGCACGATCATGCGCACCC
GTGGGGCCGCCATGCCGGCGATAATGGCCTGCTTCTCGCCGAAACGTTTGGTGGCGGGACCAGTGA
CGAAGGCTTGAGCGAGGGCGTGCAAGATTCCGAATACCGCAAGCGACAGGCCGATCATCGTCGCGC
TCCAGCGAAAGCGGTCCTCGCCGAAAATGACCCAGAGCGCTGCCGGCACCTGTCCTACGAGTTGCAT
GATAAAGAAGACAGTCATAAGTGCGGCGACGATAGTCATGCCCCGCGCCCACCGGAAGGAGCTGAC

TGGGTTGAAGGCTCTCAAGGGCATCGGTCGAGATCCCGGTGCCTAATGAGTGAGCTAACTTACATTA
ATTGCGTTGCGCTCACTGCCCGCTTTCCAGTCGGGAAACCTGTCGTGCCAGCTGCATTAATGAATCGG
CCAACGCGCGGGGAGAGGCGGTTTGCGTATTGGGCGCCAGGGTGGTTTTTCTTTTCACCAGTGAGA
CGGGCAACAGCTGATTGCCCTTCACCGCCTGGCCCTGAGAGAGTTGCAGCAAGCGGTCCACGCTGGT
TTGCCCCAGCAGGCGAAAATCCTGTTTGATGGTGGTTAACGGCGGGATATAACATGAGCTGTCTTCG
GTATCGTCGTATCCCACTACCGAGATATCCGCACCAACGCGCAGCCCGGACTCGGTAATGGCGCGCA
TTGCGCCCAGCGCCATCTGATCGTTGGCAACCAGCATCGCAGTGGGAACGATGCCCTCATTCAGCAT
TTGCATGGTTTGTTGAAAACCGGACATGGCACTCCAGTCGCCTTCCCGTTCCGCTATCGGCTGAATTT
GATTGCGAGTGAGATATTTATGCCAGCCAGCCAGACGCAGACGCGCCGAGACAGAACTTAATGGGC
CCGCTAACAGCGCGATTTGCTGGTGACCCAATGCGACCAGATGCTCCACGCCCAGTCGCGTACCGTC
TTCATGGGAGAAAATAATACTGTTGATGGGTGTCTGGTCAGAGACATCAAGAAATAACGCCGGAAC
ATTAGTGCAGGCAGCTTCCACAGCAATGGCATCCTGGTCATCCAGCGGATAGTTAATGATCAGCCCA
CTGACGCGTTGCGCGAGAAGATTGTGCACCGCCGCTTTACAGGCTTCGACGCCGCTTCGTTCTACCAT
CGACACCACCACGCTGGCACCCAGTTGATCGGCGCGAGATTTAATCGCCGCGACAATTTGCGACGGC
GCGTGCAGGGCCAGACTGGAGGTGGCAACGCCAATCAGCAACGACTGTTTGCCCGCCAGTTGTTGT
GCCACGCGGTTGGGAATGTAATTCAGCTCCGCCATCGCCGCTTCCACTTTTTCCCGCGTTTTCGCAGA
AACGTGGCTGGCCTGGTTCACCACGCGGGAAACGGTCTGATAAGAGACACCGGCATACTCTGCGAC
ATCGTATAACGTTACTGGTTTCACATTCACCACCCTGAATTGACTCTCTTCCGGGCGCTATCATGCCAT
ACCGCGAAAGGTTTTGCGCCATTCGATGGTGTCCGGGATCTCGACGCTCTCCCTTATGCGACTCCTGC
ATTAGGAAGCAGCCCAGTAGTAGGTTGAGGCCGTTGAGCACCGCCGCCGCAAGGAATGGTGCATGC
AAGGAGATGGCGCCCAACAGTCCCCCGGCCACGGGGCCTGCCACCATACCCACGCCGAAACAAGCG
CTCATGAGCCCGAAGTGGCGAGCCCGATCTTCCCCATCGGTGATGTCGGCGATATAGGCGCCAGCAA
CCGCACCTGTGGCGCCGGTGATGCCGGCCACGATGCGTCCGGCGTAGAGGATCGAGATCTCGATCC
CGCGAAATTAATACGACTCACTATAGGGGAATTGTGAGCGGATAACAATTCCCCTCTAGAAATAATT
TTGTTTAACTTTAAGAAGGAGATATACCATGAAAAAGACAGCTATCGCGATTGCAGTGGCACTGGCT
GGTTTCGCTACCGTAGCGCAGGCCGCTCCGCAAGATAACACTAGCGCCATGGGCAAACATCACCATC
ACCATCACCCCATGAGCGATTACGACATCCCCACTACTAAGCTTCTGGAAGTTCTGTTCCAGGGGCCC
CATATGACTGGCTGTGTATAAGGGAGCCTGACATTTATATTCCCCAGAACATCAGGTTAATGGCGTTT
TTGATGTCATTTTCGCGGTGGCTGAGATCAGCCACTTCTTCCCCGATAACGGAGACCGGCACACTGG
CCATATCGGTGGTCATCATGCGCCAGCTTTCATCCCCGATATGCACCACCGGGTAAAGTTCACGGGA
GACTTTATCTGACAGCAGACGTGCACTGGCCAGGGGGATCACCATCCGTCGCCCGGGCGTGTCAATA
ATATCACTCTGTACATCCACAAACAGACGATAACGGCTCTCTCTTTTATAGGTGTAAACCTTAAACTGC
ATTTCACCAGCCCCTGTTCTCGTCAGCAAAAGAGCCGTTCATTTCAATAAACCGGGCGACCTCAGCCA
TCCCTTCCTGATTTTCCGCTTTCCAGCGTTCGGCACGCAGACGACGGGCTTCATTCTGCATGGTTGTG
CTTACCAGACCGGAGATATTGACATCATATATGCCTTGAGCAACTGATAGCTGTCGCTGTCAACTGTC
ACTGTAATACGCTGCTTCATAGCATACCTCTTTTTGACATACTTCGGGTATACATATCAGTATATATTC
TTATACCGCAAAAATCAGCGCGCAAATACGCATACTGTTATCTGGCTTTTAGTAAGCCGGATCCACGC
GTCTCGAGCACCACCACCACCACCACTGAGATCCGGCTGCTAACAAAGCCCGAAAGGAAGCTGAGTT
GGCTGCTGCCACCGCTGAGCAATAACTAGCATAACCCCTTGGGGCCTCTAAACGGGTCTTGAGGGGT
TTTTTGCTGAAAGGAGGAACTATATCCGGAT

> pDB_ccdb_his_MBP_3C
TGGCGAATGGGACGCGCCCTGTAGCGGCGCATTAAGCGCGGCGGGTGTGGTGGTTACGCGCAGCG
TGACCGCTACACTTGCCAGCGCCCTAGCGCCCGCTCCTTTCGCTTTCTTCCCTTCCTTTCTCGCCACGTT
CGCCGGCTTTCCCCGTCAAGCTCTAAATCGGGGGCTCCCTTTAGGGTTCCGATTTAGTGCTTTACGGC
ACCTCGACCCCAAAAAACTTGATTAGGGTGATGGTTCACGTAGTGGGCCATCGCCCTGATAGACGGT
TTTTCGCCCTTTGACGTTGGAGTCCACGTTCTTTAATAGTGGACTCTTGTTCCAAACTGGAACAACACT

CAACCCTATCTCGGTCTATTCTTTTGATTTATAAGGGATTTTGCCGATTTCGGCCTATTGGTTAAAAAA
TGAGCTGATTTAACAAAAATTTAACGCGAATTTTAACAAAATATTAACGTTTACAATTTCAGGTGGCA
CTTTTCGGGGAAATGTGCGCGGAACCCCTATTTGTTTATTTTTCTAAATACATTCAAATATGTATCCGC
TCATGAATTAATTCTTAGAAAAACTCATCGAGCATCAAATGAAACTGCAATTTATTCATATCAGGATT
ATCAATACCATATTTTTGAAAAAGCCGTTTCTGTAATGAAGGAGAAAACTCACCGAGGCAGTTCCATA
GGATGGCAAGATCCTGGTATCGGTCTGCGATTCCGACTCGTCCAACATCAATACAACCTATTAATTTC
CCCTCGTCAAAAATAAGGTTATCAAGTGAGAAATCACCATGAGTGACGACTGAATCCGGTGAGAATG
GCAAAAGTTTATGCATTTCTTTCCAGACTTGTTCAACAGGCCAGCCATTACGCTCGTCATCAAAATCAC
TCGCATCAACCAAACCGTTATTCATTCGTGATTGCGCCTGAGCGAGACGAAATACGCGATCGCTGTTA
AAAGGACAATTACAAACAGGAATCGAATGCAACCGGCGCAGGAACACTGCCAGCGCATCAACAATA
TTTTCACCTGAATCAGGATATTCTTCTAATACCTGGAATGCTGTTTTCCCGGGGATCGCAGTGGTGAG
TAACCATGCATCATCAGGAGTACGGATAAAATGCTTGATGGTCGGAAGAGGCATAAATTCCGTCAGC
CAGTTTAGTCTGACCATCTCATCTGTAACATCATTGGCAACGCTACCTTTGCCATGTTTCAGAAACAAC
TCTGGCGCATCGGGCTTCCCATACAATCGATAGATTGTCGCACCTGATTGCCCGACATTATCGCGAGC
CCATTTATACCCATATAAATCAGCATCCATGTTGGAATTTAATCGCGGCCTAGAGCAAGACGTTTCCC
GTTGAATATGGCTCATAACACCCCTTGTATTACTGTTTATGTAAGCAGACAGTTTTATTGTTCATGACC
AAAATCCCTTAACGTGAGTTTTCGTTCCACTGAGCGTCAGACCCCGTAGAAAAGATCAAAGGATCTTC
TTGAGATCCTTTTTTTCTGCGCGTAATCTGCTGCTTGCAAACAAAAAAACCACCGCTACCAGCGGTGG
TTTGTTTGCCGGATCAAGAGCTACCAACTCTTTTTCCGAAGGTAACTGGCTTCAGCAGAGCGCAGATA
CCAAATACTGTCCTTCTAGTGTAGCCGTAGTTAGGCCACCACTTCAAGAACTCTGTAGCACCGCCTAC
ATACCTCGCTCTGCTAATCCTGTTACCAGTGGCTGCTGCCAGTGGCGATAAGTCGTGTCTTACCGGGT
TGGACTCAAGACGATAGTTACCGGATAAGGCGCAGCGGTCGGGCTGAACGGGGGGTTCGTGCACA
CAGCCCAGCTTGGAGCGAACGACCTACACCGAACTGAGATACCTACAGCGTGAGCTATGAGAAAGC
GCCACGCTTCCCGAAGGGAGAAAGGCGGACAGGTATCCGGTAAGCGGCAGGGTCGGAACAGGAGA
GCGCACGAGGGAGCTTCCAGGGGGAAACGCCTGGTATCTTTATAGTCCTGTCGGGTTTCGCCACCTC
TGACTTGAGCGTCGATTTTTGTGATGCTCGTCAGGGGGGCGGAGCCTATGGAAAAACGCCAGCAAC
GCGGCCTTTTTACGGTTCCTGGCCTTTTGCTGGCCTTTTGCTCACATGTTCTTTCCTGCGTTATCCCCTG
ATTCTGTGGATAACCGTATTACCGCCTTTGAGTGAGCTGATACCGCTCGCCGCAGCCGAACGACCGA
GCGCAGCGAGTCAGTGAGCGAGGAAGCGGAAGAGCGCCTGATGCGGTATTTTCTCCTTACGCATCT
GTGCGGTATTTCACACCGCATATATGGTGCACTCTCAGTACAATCTGCTCTGATGCCGCATAGTTAAG
CCAGTATACACTCCGCTATCGCTACGTGACTGGGTCATGGCTGCGCCCCGACACCCGCCAACACCCG
CTGACGCGCCCTGACGGGCTTGTCTGCTCCCGGCATCCGCTTACAGACAAGCTGTGACCGTCTCCGG
GAGCTGCATGTGTCAGAGGTTTTCACCGTCATCACCGAAACGCGCGAGGCAGCTGCGGTAAAGCTC
ATCAGCGTGGTCGTGAAGCGATTCACAGATGTCTGCCTGTTCATCCGCGTCCAGCTCGTTGAGTTTCT
CCAGAAGCGTTAATGTCTGGCTTCTGATAAAGCGGGCCATGTTAAGGGCGGTTTTTTCCTGTTTGGTC
ACTGATGCCTCCGTGTAAGGGGGATTTCTGTTCATGGGGGTAATGATACCGATGAAACGAGAGAGG
ATGCTCACGATACGGGTTACTGATGATGAACATGCCCGGTTACTGGAACGTTGTGAGGGTAAACAAC
TGGCGGTATGGATGCGGCGGGACCAGAGAAAAATCACTCAGGGTCAATGCCAGCGCTTCGTTAATA
CAGATGTAGGTGTTCCACAGGGTAGCCAGCAGCATCCTGCGATGCAGATCCGGAACATAATGGTGC
AGGGCGCTGACTTCCGCGTTTCCAGACTTTACGAAACACGGAAACCGAAGACCATTCATGTTGTTGC
TCAGGTCGCAGACGTTTTGCAGCAGCAGTCGCTTCACGTTCGCTCGCGTATCGGTGATTCATTCTGCT
AACCAGTAAGGCAACCCCGCCAGCCTAGCCGGGTCCTCAACGACAGGAGCACGATCATGCGCACCC
GTGGGGCCGCCATGCCGGCGATAATGGCCTGCTTCTCGCCGAAACGTTTGGTGGCGGGACCAGTGA
CGAAGGCTTGAGCGAGGGCGTGCAAGATTCCGAATACCGCAAGCGACAGGCCGATCATCGTCGCGC
TCCAGCGAAAGCGGTCCTCGCCGAAAATGACCCAGAGCGCTGCCGGCACCTGTCCTACGAGTTGCAT
GATAAAGAAGACAGTCATAAGTGCGGCGACGATAGTCATGCCCCGCGCCCACCGGAAGGAGCTGAC
TGGGTTGAAGGCTCTCAAGGGCATCGGTCGAGATCCCGGTGCCTAATGAGTGAGCTAACTTACATTA

ATTGCGTTGCGCTCACTGCCCGCTTTCCAGTCGGGAAACCTGTCGTGCCAGCTGCATTAATGAATCGG
CCAACGCGCGGGGAGAGGCGGTTTGCGTATTGGGCGCCAGGGTGGTTTTTCTTTTCACCAGTGAGA
CGGGCAACAGCTGATTGCCCTTCACCGCCTGGCCCTGAGAGAGTTGCAGCAAGCGGTCCACGCTGGT
TTGCCCCAGCAGGCGAAAATCCTGTTTGATGGTGGTTAACGGCGGGATATAACATGAGCTGTCTTCG
GTATCGTCGTATCCCACTACCGAGATATCCGCACCAACGCGCAGCCCGGACTCGGTAATGGCGCGCA
TTGCGCCCAGCGCCATCTGATCGTTGGCAACCAGCATCGCAGTGGGAACGATGCCCTCATTCAGCAT
TTGCATGGTTTGTTGAAAACCGGACATGGCACTCCAGTCGCCTTCCCGTTCCGCTATCGGCTGAATTT
GATTGCGAGTGAGATATTTATGCCAGCCAGCCAGACGCAGACGCGCCGAGACAGAACTTAATGGGC
CCGCTAACAGCGCGATTTGCTGGTGACCCAATGCGACCAGATGCTCCACGCCCAGTCGCGTACCGTC
TTCATGGGAGAAAATAATACTGTTGATGGGTGTCTGGTCAGAGACATCAAGAAATAACGCCGGAAC
ATTAGTGCAGGCAGCTTCCACAGCAATGGCATCCTGGTCATCCAGCGGATAGTTAATGATCAGCCCA
CTGACGCGTTGCGCGAGAAGATTGTGCACCGCCGCTTTACAGGCTTCGACGCCGCTTCGTTCTACCAT
CGACACCACCACGCTGGCACCCAGTTGATCGGCGCGAGATTTAATCGCCGCGACAATTTGCGACGGC
GCGTGCAGGGCCAGACTGGAGGTGGCAACGCCAATCAGCAACGACTGTTTGCCCGCCAGTTGTTGT
GCCACGCGGTTGGGAATGTAATTCAGCTCCGCCATCGCCGCTTCCACTTTTTCCCGCGTTTTCGCAGA
AACGTGGCTGGCCTGGTTCACCACGCGGGAAACGGTCTGATAAGAGACACCGGCATACTCTGCGAC
ATCGTATAACGTTACTGGTTTCACATTCACCACCCTGAATTGACTCTCTTCCGGGCGCTATCATGCCAT
ACCGCGAAAGGTTTTGCGCCATTCGATGGTGTCCGGGATCTCGACGCTCTCCCTTATGCGACTCCTGC
ATTAGGAAGCAGCCCAGTAGTAGGTTGAGGCCGTTGAGCACCGCCGCCGCAAGGAATGGTGCATGC
AAGGAGATGGCGCCCAACAGTCCCCCGGCCACGGGGCCTGCCACCATACCCACGCCGAAACAAGCG
CTCATGAGCCCGAAGTGGCGAGCCCGATCTTCCCCATCGGTGATGTCGGCGATATAGGCGCCAGCAA
CCGCACCTGTGGCGCCGGTGATGCCGGCCACGATGCGTCCGGCGTAGAGGATCGAGATCTCGATCC
CGCGAAATTAATACGACTCACTATAGGGGAATTGTGAGCGGATAACAATTCCCCTCTAGAAATAATT
TTGTTTAACTTTAAGAAGGAGATATACCATGGGCAGCAGCCATCATCATCATCATCACGGTACCAAAA
CTGAAGAAGGTAAACTGGTAATCTGGATTAACGGCGATAAAGGCTATAACGGTCTCGCTGAAGTCG
GTAAGAAATTCGAGAAAGATACCGGAATTAAAGTCACCGTTGAGCATCCGGATAAACTGGAAGAGA
AATTCCCACAGGTTGCGGCAACTGGCGATGGCCCTGACATTATCTTCTGGGCACACGACCGCTTTGG
TGGCTACGCTCAATCTGGCCTGTTGGCTGAAATCACCCCGGACAAAGCGTTCCAGGACAAGCTGTAT
CCGTTTACCTGGGATGCCGTACGTTACAACGGCAAGCTGATTGCTTACCCGATCGCTGTTGAAGCGTT
ATCGCTGATTTATAACAAAGATCTGCTGCCGAACCCGCCAAAAACCTGGGAAGAGATCCCGGCGCTG
GATAAAGAACTGAAAGCGAAAGGTAAGAGCGCGCTGATGTTCAACCTGCAAGAACCGTACTTCACC
TGGCCGCTGATTGCTGCTGACGGGGGTTATGCGTTCAAGTATGAAAACGGCAAGTACGACATTAAA
GACGTGGGCGTGGATAACGCTGGCGCGAAAGCGGGTCTGACCTTCCTGGTTGACCTGATTAAAAAC
AAACACATGAATGCAGACACCGATTACTCCATCGCAGAAGCTGCCTTTAATAAAGGCGAAACAGCGA
TGACCATCAACGGCCCGTGGGCATGGTCCAACATCGACACCAGCAAAGTGAATTATGGTGTAACGGT
ACTGCCGACCTTCAAGGGTCAACCATCCAAACCGTTCGTTGGCGTGCTGAGCGCAGGTATTAACGCC
GCCAGTCCGAACAAAGAGCTGGCGAAAGAGTTCCTCGAAAACTATCTGCTGACTGATGAAGGTCTG
GAAGCGGTTAATAAAGACAAACCGCTGGGTGCCGTAGCGCTGAAGTCTTACGAGGAAGAGTTGGCG
AAAGATCCACGTATTGCCGCCACCATGGAAAACGCCCAGAAAGGTGAAATCATGCCGAACATCCCGC
AGATGTCCGCTTTCTGGTATGCCGTGCGTACTGCGGTGATCAACGCCGCCAGCGGTCGTCAGACTGT
CGATGAAGCCCTGAAAGACGCGCAGACTGGTACCGATTACGATATCCCAACGACCAAGCTTCTGGAA
GTTCTGTTCCAGGGGCCCCATATGACTGGCTGTGTATAAGGGAGCCTGACATTTATATTCCCCAGAAC
ATCAGGTTAATGGCGTTTTTGATGTCATTTTCGCGGTGGCTGAGATCAGCCACTTCTTCCCCGATAAC
GGAGACCGGCACACTGGCCATATCGGTGGTCATCATGCGCCAGCTTTCATCCCCGATATGCACCACC
GGGTAAAGTTCACGGGAGACTTTATCTGACAGCAGACGTGCACTGGCCAGGGGGATCACCATCCGT
CGCCCGGGCGTGTCAATAATATCACTCTGTACATCCACAAACAGACGATAACGGCTCTCTCTTTTATA
GGTGTAAACCTTAAACTGCATTTCACCAGCCCCTGTTCTCGTCAGCAAAAGAGCCGTTCATTTCAATA

AACCGGGCGACCTCAGCCATCCCTTCCTGATTTTCCGCTTTCCAGCGTTCGGCACGCAGACGACGGGC
TTCATTCTGCATGGTTGTGCTTACCAGACCGGAGATATTGACATCATATATGCCTTGAGCAACTGATA
GCTGTCGCTGTCAACTGTCACTGTAATACGCTGCTTCATAGCATACCTCTTTTTGACATACTTCGGGTA
TACATATCAGTATATATTCTTATACCGCAAAAATCAGCGCGCAAATACGCATACTGTTATCTGGCTTTT
AGTAAGCCGGATCCACGCGTCTCGAGCACCACCACCACCACCACTGAGATCCGGCTGCTAACAAAGC
CCGAAAGGAAGCTGAGTTGGCTGCTGCCACCGCTGAGCAATAACTAGCATAACCCCTTGGGGCCTCT
AAACGGGTCTTGAGGGGTTTTTTGCTGAAAGGAGGAACTATATCCGGAT

**Supplemental table 1: Summary of QtPISA analysis of the minimized X-ray and model structures of MAX/HMA complexes**

| Complex | BE cal.mol$^{-1}$ | ΔiG kcal.mol$^{-1}$ | Total interface area Å$^2$ | No. of H-Bonds | No. of Salt Bridges |
|---|---|---|---|---|---|
| **AVR1-CO39/RGA5_HMA** (PDB 5ZNG) | -6.5(-7.3) | -3.0(-4.6) | 499.0(492.8) | 7(6) | 1(0) |
| **AVR-Pia/Pikp-1_HMA** (PDB 6Q76) | -8.9(-8.6) | -4.6(-4.7) | 491.6(460.7) | 7(7) | 3(2) |
| **AVR-PiKD/Pikp-1_HMA** (PDB 6G10) | -10.3(-10.8) | -2.1(-2.2) | 1106.6(979.3) | 11(10) | 9(11) |
| **AVR-Pia/RGA5_HMA** | -7.2 | -4.3 | 488.0 | 4 | 3 |
| **AVR-Pia/HMAm1m2** | -8.6 | -4.3 | 481.4 | 8 | 2 |
| **AVR1-CO39/HMAm1m2** | -6.8 | -3.4 | 482.8 | 6 | 2 |
| **AVR-PikD/HMAm1m2** | -11.3 | -2.2 | 1052.1 | 13 | 9 |

As controls, the three PDB structures (5ZNG, 6Q76 and 6G10) used as templates for modelling HMA domains in complex with MAX effectors were minimized with Charmm and analysed with QtPISA. The values given between parenthesis correspond to the initial PDB structures without minimization. In the bottom section, five models of MAX/HMA complexes generated as described in the Methods were refined with Charmm and analysed with QtPISA. BE is the total Binding Energy at the interaction interface and ΔiG is the solvation energy gain. The number of H-bonds and salt-bridges formed at the complex interface are also indicated.

**Supplemental Table 2: Binding and fitting parameters of AVR-PikD interaction with different HMA domains, calculated for the kinetic titrations shown in supplementary Figure 4 using the indicated interaction model for fitting SPR data.**

| **Steady-state model** | **$K_D$ (M)** | $R_{max}$ (RU) | Chi² (RU²) |
|---|---|---|---|
| AVR-PikD/RGA5_HMA wt | **8.32E-06** | 861 | 9.51 |

| **Heterogeneous model** | **$K_D$1 (M)** | $k_a$1 (1/Ms) | $k_d$1 (1/s) | $K_D$2 (M) | $k_a$2 (1/Ms) | $k_d$2 (1/s) | $R_{max}$1 (RU) | $R_{max}$2 (RU) | Chi² (RU²) |
|---|---|---|---|---|---|---|---|---|---|
| AVR-PikD/RGA5_HMAm1 | **1.60E-09** | 7.30E+05 | 1.17E-03 | 2.90E-07 | 6.66E+03 | 1.93E-03 | 598 | 521 | 29.4 |
| AVR-PikD/RGA5_HMAm1m2 | **5.38E-10** | 8.36E+05 | 4.50E-04 | 3.60E-07 | 8.61E+03 | 3.10E-03 | 549 | 517 | 3.98 |
| AVR-PikD/Pikp-1_HMA | **8.30E-10** | 1.43E+06 | 1.19E-03 | 4.44E-08 | 3.84E+05 | 1.71E-02 | 372 | 119 | 11.9 |

| Genes transformed | Independant transgenic line | Genotype RGA4 (T0) | Genotype RGA5 (T0) | Sequencing RGA5 PCR products (T0) | Phenotype T0 Guy11_EV | Phenotype T0 Guy11_AVR-Pia | Phenotype T0 JP10 (AVR-PikD+) | T0 seed production | Phenotype T1 Guy11_EV | Phenotype T1 Guy11_AVR-Pia | Phenotype T1 Guy11_AVR1-CO39 | Phenotype T1 Guy11_AVR-PikD | Phenotype T2 Guy11_AVR-Pia | Phenotype T2 Guy11_AVR1-CO39 | Phenotype T2 Guy11_AVR-PikD |
|---|---|---|---|---|---|---|---|---|---|---|---|---|---|---|---|
| RGA4+RGA5 | L1 | + | + | + | S | S | S | 20 | nt | nt | nt | nt | nt | nt | nt |
| | L2 | + | + | nt | S | R | S | >100 | S | R | R | S | R | R | S |
| | L3 | + | + | + | nt | nt | nt | ~50 | nt | nt | nt | nt | nt | nt | nt |
| | lacking RGA5 | + | - | nt | nt | nt | nt | >100 | nt | nt | nt | nt | nt | nt | nt |
| | L4 | + | + | + | S | nt | S | 5 | nt | nt | nt | nt | nt | nt | nt |
| | L5 | + | + | + | S | R | S | ~50 | S | R | R | S | nt | nt | nt |
| | L6 | + | + | nt | S | nt | S | >100 | nt | nt | nt | nt | nt | nt | nt |
| | L7 | + | + | + | nt | nt | nt | 10 | nt | nt | nt | nt | nt | nt | nt |
| | lacking RGA5 | + | - | nt | nt | nt | S | >100 | nt | nt | nt | nt | nt | nt | nt |
| | L8 | + | + | nt | S | S | S | >100 | nt | nt | nt | nt | nt | nt | nt |
| | L9 | + | + | nt | nt | R | nt | 0 | nt | nt | nt | nt | nt | nt | nt |
| | L10 | + | + | nt | S | R | nt | 0 | nt | nt | nt | nt | nt | nt | nt |
| | L11 | + | + | nt | nt | R | nt | 0 | nt | nt | nt | nt | nt | nt | nt |
| | L12 | + | + | nt | S | R | nt | >100 | S | R | R | S | R | R | S |
| | L13 | + | + | nt | nt | nt | nt | 0 | nt | nt | nt | nt | nt | nt | nt |
| | L14 | + | + | nt | S | R | S | 10 | nt | nt | nt | nt | nt | nt | nt |
| RGA4+RGA5m1 | L1 | + | + | + | S | nt | S | >100 | S | R | R | S | nt | nt | nt |
| | L2 | + | + | + | S | S | S | >100 | nt | nt | nt | nt | nt | nt | nt |
| | L3 | + | + | + | S | R | S | ~50 | S | R | R | S | R | R | S |
| | L4 | + | + | + | nt | nt | nt | 0 | nt | nt | nt | nt | nt | nt | nt |
| | L5 | + | + | + | nt | nt | nt | 0 | nt | nt | nt | nt | nt | nt | nt |
| | L6 | + | + | + | nt | nt | nt | 0 | nt | nt | nt | nt | nt | nt | nt |
| RGA4+RGA5m2 | L1 | + | + | + | S | R | S | >100 | S | R | R | S | R | R | S |
| | L2 | + | + | + | nt | nt | nt | >100 | nt | nt | nt | nt | R | R | S |
| | L3 | + | + | + | S | R | S | 9 | nt | nt | nt | nt | nt | nt | nt |
| | L4 | + | + | + | nt | nt | nt | >100 | S | R | R | S | R | R | S |
| | L5 | + | + | + | S | R | S | ~50 | S | R | 1 dwarf plant | S | R | R | S |
| RGA4+RGA5m1m2 | L1 | + | + | + | S | R | S | 0 | nt | nt | nt | nt | nt | nt | nt |
| | L2 | + | + | + | nt | nt | nt | 0 | nt | nt | nt | nt | nt | nt | nt |
| | L3 | + | + | + | S | R | S | ~50 | S | R | R | S | R | R | S |
| | L4 | + | + | + | S | S | S | ~50 | nt | nt | nt | nt | nt | nt | nt |
| | L5 | + | + | + | nt | R | S | 4 | nt | nt | nt | nt | R | R | S |
| | L6 | + | + | + | S | S | S | ~50 | nt | nt | nt | nt | nt | nt | nt |
| RGA4+GFP | L1 | + | - | nt | nt | nt | nt | 10 | nt | nt | nt | nt | nt | nt | nt |
| | L2 | + | - | nt | nt | nt | nt | 10 | nt | nt | nt | nt | nt | nt | nt |
| | L3 | + | - | nt | nt | nt | nt | 10 | nt | nt | nt | nt | nt | nt | nt |
| | L4 | + | - | nt | nt | nt | nt | 0 | nt | nt | nt | nt | nt | nt | nt |
| | L5 | + | - | nt | nt | nt | nt | 0 | nt | nt | nt | nt | nt | nt | nt |
| | L6 | + | - | nt | S | S | S | 20 | nt | nt | nt | nt | nt | nt | nt |
| | L7 | + | - | nt | nt | nt | nt | 10 | nt | nt | nt | nt | nt | nt | nt |
| | L8 | + | - | nt | S | S | S | >100 | nt | nt | nt | nt | nt | nt | nt |
| | L9 | + | - | nt | nt | nt | nt | 0 | nt | nt | nt | nt | nt | nt | nt |
| | L10 | + | - | nt | S | S | S | >100 | nt | nt | nt | nt | nt | nt | nt |
| | L11 | + | - | nt | S | S | S | 4 | nt | nt | nt | nt | nt | nt | nt |
| | lacking RGA4 | - | - | nt | nt | nt | nt | >100 | nt | nt | nt | nt | nt | nt | nt |
| | L12 | + | - | nt | S | S | S | >100 | S | S | S | S | S | S | S |
| | lacking RGA4 | - | - | nt | S | S | S | >100 | nt | nt | nt | nt | nt | nt | nt |
| | L13 | + | - | nt | nt | nt | nt | >100 | S | S | S | S | S | S | S |
| | L14 | + | - | nt | nt | nt | nt | >100 | S | S | S | S | nt | nt | nt |
| / | / | / | / | / | I | I | R | / | S | S | S | R | S | S | R |
| / | / | / | / | / | S | S | S | / | S | S | S | S | S | S | S |
| / | / | / | / | / | S | S | S | / | S | S | S | S | S | S | S |
| / | / | / | / | / | S | S | R | / | nt | nt | nt | nt | nt | nt | nt |
| / | / | / | / | / | S | S | R | / | nt | nt | nt | nt | nt | nt | nt |
| / | / | / | / | / | S | R | R | / | nt | nt | nt | nt | nt | nt | nt |

**Supplemental Table 3: Inoculation experiments on T0, T1 and T2 rice transgenic plants.** S = susceptible; R = resistant; I = intermediate resistance; nt = not tested.

**Supplemental Table 4: Primers**

| Primers | Sequence |
|---|---|
| oCS080 | GGGGACCACTTTGTACAAGAAAGCTGGGTCTCACATGGTTGAGCAAGGGTTAA |
| oCS108 | GGGGACAAGTTTGTACAAAAAAGCAGGCTTAATGGGTGGCGGGTGGACTAACAA |
| oCS109 | GGGGACCACTTTGTACAAGAAAGCTGGGTCTTACATAATATTGCAGCCCTCTTC |
| oCS118 | GGGGACAAGTTTGTACAAAAAAGCAGGCTTAATGGAGCACCTTGTAAGCCTCCAG |
| oCS209 | GGGGACAAGTTTGTACAAAAAAGCAGGCTTAATGGTGAGCAAGGGCGAGG |
| oCS210 | GGGGACCACTTTGTACAAGAAAGCTGGGTCTTATAAGCCTGCTTTTTTGTACAAACTTG |
| oCS330 | GTGAACGGGGTGGACAGCGTGGCGTTAGTGGGGGATCTAAGAGACAAGATCGAGGTGGTCGGCCGTGGCATTGAC |
| oCS331 | GTCAATGCCACGGCCGACCACCTCGATCTTGTCTCTTAGATCCCCCACTAACGCCACGCTGTCCACCCCGTTCAC |
| oCS332 | GAAATGTGGCCTCGCCGAGCTCTTGCAGGTGTCGCAGGTTGAGAAAGAGAAGACACAGCTGG |
| oCS333 | CCAGCTGTGTCTTCTCTTTCTCAACCTGCGACACCTGCAAGAGCTCGGCGAGGCCACATTTC |
| oCS334 | GGGGACAAGTTTGTACAAAAAAGCAGGCTTAATGGAAACGGGCAACAAATATATAGAAAAA |
| oCS335 | GGGGACCACTTTGTACAAGAAAGCTGGGTCTTAAAAGCCGGGCCTTTTTTTC |
| oCS346 | GGGGACAAGTTTGTACAAAAAAGCAGGCTTACTCAGAAAAACAGGGCTAAAGCAAAA |
| oCS347 | GGGGACCACTTTGTACAAGAAAGCTGGGTCTCAATCTTTATTTGCTTGGCTGACCTGC |
| oCS348 | GGGGACAAGTTTGTACAAAAAAGCAGGCTTAAGTGCATTAACGGGGCAACG |
| oCS349 | GGGGACCACTTTGTACAAGAAAGCTGGGTCTCACTCTTTCTCAACTAACTCCACCATCAA |
| oCS351 | GGGGACCACTTTGTACAAGAAAGCTGGGTCTCACTCTTTCTCAACCTGCGACACCT |
| oCS385 | GCTTGCAGGGGTGGACAGCGTGGCGTTAGTGGGGGATCTAAGAGACAAGATCGAGGTGGTCGGCCGTGGCATTGAC |
| oCS386 | GTCAATGCCACGGCCGACCACCTCGATCTTGTCTCTTAGATCCCCCACTAACGCCACGCTGTCCACCCCTGCAAGC |
| oTK017_c | AGGACCCAATCTTCAAAATGGCTAAAGTCCAAGCTACATTCGCCA |
| oTK036 | GGAGCCTGAATGTTGAGTGGAATGATGCGGGATCAACAAGACTCATCGTCGTCA |
| oTK439 | TATCATATGGCTGCGCCAGCTAGATCTTGCGTCTAT |
| oTK334 | TATGGATCCCTAGTAAGGCTCGGCAGCAAG |

### Supplemental Table 5: Constructs.

| Use | Final plasmid | Cloning Method | Template | Cloning primers | Backbone vector | Insert | Reference |
|---|---|---|---|---|---|---|---|
| Entry vectors for LR cloning | pSC049 | / | / | / | pDONR207 | dSP-AVR-Pia with start without stop (20-85) | Cesari et al., 2013 |
| | pSC060 | / | / | / | pDONR207 | dSP-AVR-Pia with start with stop (20-85) | Cesari et al., 2014 |
| | pSC120 | GTW BP | pTH123.1 | oCS108/oCS109 | pDONR207 | dSP-PWL2 with start with stop (22-145) | |
| | pSC129 | / | / | / | pDONR207 | RGA5_Cter with stop (883-1116) | Cesari et al., 2013 |
| | pSC210 | / | / | / | pDONR207 | RGA5_deltaHMA with stop (1-996) | Cesari et al., 2014 |
| | pSC309 | GTW BP | pBIN19-YFP-GTW | oCS209/oCS210 | pDONR207 | YFP with stop | Cesari et al., 2014 |
| | pSC467 | Quikchange | pSC057 | oCS330/oCS331 | pDONR207 | RGA5m1 with stop (1-1116) | Cesari et al., 2014 |
| | pSC468 | quikchange | pSC057 | oCS332/oCS333 | pDONR207 | RGA5m2 with stop (1-1116) | |
| | pSC469 | quikchange | pSC467 | oCS332/oCS333 | pDONR207 | RGA5m1m2 with stop (1-1116) | |
| | pSC470 | quikchange | pSC207 | oCS330/oCS331 | pDONR207 | RGA5_HMAm1 with stop (997-1072) | |
| | pSC472 | GTW BP | pSC469 | oCS348/oCS351 | pDONR207 | RGA5_HMAm1m2 with stop (991-1072) | |
| | pSC474 | GTW-BP | pTK210 | oCS334/oCS335 | pDONR207 | dSP-AVR-PikD with stop (22-113) | |
| | pSC500 | GTW-BP | pSC496 | oCS346/oCS347 | pDONR207 | Pikp-1_HMA with stop (182-263) | |
| | pSC501 | GTW-BP | pSC057 | oCS348/oCS349 | pDONR207 | RGA5_HMA with stop (991-1072) | |
| | pSC502 | GTW-BP | pSC469 | oCS118/oCS080 | pDONR207 | RGA5_C-ter_m1m2 with stop (883-1116) | |
| | pSC629 | GTW-BP | pSC467 | oCS348/oCS349 | pDONR207 | RGA5_HMAm1 with stop (991-1072) | |
| | pSC630 | GTW-BP | pSC468 | oCS348/oCS351 | pDONR207 | RGA5_HMAm2 with stop (991-1072) | |
| | pSC631 | GTW-BP | pSC467 | oCS118/oCS080 | pDONR207 | RGA5_Cter_m1 with stop (883-1116) | |
| | pSC632 | GTW-BP | pSC468 | oCS118/oCS080 | pDONR207 | RGA5_Cter_m2 with stop (883-1116) | |
| Y2H | pSC001 | / | / | / | pGBKT7-BD | BD:dSP-AVR1-CO39 (22-89) | Cesari et al., 2013 |
| | pCV243 | GTW LR | pSC060 | / | pGBKT7-BD_GTW | BD:dSP-AVR-Pia (20-85) | |
| | pSC490 | GTW LR | pSC474 | / | pGBKT7-BD_GTW | BD:dSP-AVR-PikD (22-113) | |
| | pGBKT7-BD | GTW LR | / | / | pGBKT7-BD | BD | Clontech |
| | pSC548 | GTW LR | pSC501 | / | pGADT7-AD_GTW | AD:RGA5_HMA (991-1072) | |
| | pSC636 | GTW LR | pSC629 | / | pGADT7-AD_GTW | AD:RGA5_HMAm1 (991-1072) | |
| | pSC637 | GTW LR | pSC630 | / | pGADT7-AD_GTW | AD:RGA5_HMAm2 (991-1072) | |
| | pSC485 | GTW LR | pSC472 | / | pGADT7-AD_GTW | AD:RGA5_HMAm1m2 (991-1072) | |
| | pCV226 | GTW LR | pSC129 | / | pGADT7-AD_GTW | AD:RGA5_Cter (883-1116) | |
| | pSC638 | GTW LR | pSC631 | / | pGADT7-AD_GTW | AD:RGA5_Cter_m1 (883-1116) | |
| | pSC639 | GTW LR | pSC632 | / | pGADT7-AD_GTW | AD:RGA5_Cter_m2 (883-1116) | |
| | pSC549 | GTW LR | pSC502 | / | pGADT7-AD_GTW | AD:RGA5_Cter_m1m2 (883-1116) | |
| | pSC547 | GTW LR | pSC500 | / | pGADT7-AD_GTW | AD:Pikp-1_HMA (182-263) | |
| | pGADT7-AD | GTW LR | / | / | pGADT7-AD | AD | Clontech |
| Co-IP and HR tests in *N. benthamiana* | pSC065.3 | GTW LR | pSC049 | / | pBIN19-GTW-3HA | dSP_AVR-Pia:3HA (20-85) | |
| | pSC495.1.1 | GTW LR | / | / | pCambia1300 | dSP_AVR-PikD:HA (22-113) | Maqbool et al., 2015 |
| | pSC598.1 | GTW LR | / | / | pCambia1300 | dSP_AVR-PikC:HA (22-113) | Maqbool et al., 2015 |
| | pDO134 | GTW LR | pSC120 | / | pBIN19-3HA-GTW | 3HA:PWL2 (22-145) | |
| | pSC545.2 | GTW LR | pSC501 | / | pBIN19-YFP-GTW | YFP:RGA5_HMA (991-1072) | |
| | pSC478.2.1 | GTW LR | pSC470 | / | pBIN19-YFP-GTW | YFP:RGA5_HMAm1 (997-1072) | |
| | pSC480.1 | GTW LR | pSC472 | / | pBIN19-YFP-GTW | YFP:RGA5_HMAm1m2 (991-1072) | |
| | pSC544.2 | GTW LR | pSC500 | / | pBIN19-YFP-GTW | YFP:Pikp-1_HMA (182-263) | |
| | pSC276 | GTW LR | pSC210 | / | pBIN19-YFP-GTW | YFP:RGA5_deltaHMA (1-996) | |
| | pSC310 | GTW LR | pSC309 | / | pBIN19-YFP-GTW | YFP | |
| | pSC078.3 | / | / | / | pBIN19-YFP-GTW | YFP:RGA5 (1-1116) | Cesari et al., 2013 |
| | pSC475 | GTW LR | pSC467 | / | pBIN19-YFP-GTW | YFP:RGA5m1 (1-1116) | |
| | pSC476 | GTW LR | pSC468 | / | pBIN19-YFP-GTW | YFP-RGA5m2 (1-1116) | |
| | pSC477.1 | GTW LR | pSC469 | / | pBIN19-YFP-GTW | YFP:RGA5m1m2 (1-1116) | |
| | pSC061 | / | / | / | pBIN19-3HA-GTW | RGA4:3HA (1-996) | Cesari et al., 2013 |
| | pSC095 | / | / | / | pBIN19-GTW-3HA | dSP_AVR-Pia (20-85) | Cesari et al., 2014 |
| | pSC497 | / | / | / | pICSL4723 | Pikp-1:Hellfire/Pikp-2:HA | provided by Mark Banfield |
| *Magnaporthe oryzae* transformation | pCV10 | yeast gap repair | genomic DNA of *M. oryzae* IN03 | oTK017_c/oTK036 | pDL02 | pRP27::SP-AVR1-CO39 (1-89) | (identical to pCB027 from Ribot et al, 2013 ) |
| | pCB1004-pex22 | / | / | / | pCB1004 | pAVR-Pia::SP-AVR-Pia (1-85) | Yoshida et al., 2009 |
| | pCB1004-pex31 | / | / | / | pCB1004 | pAVR-PikD::SP-AVR-PikD (1-113) | Yoshida et al., 2009 |
| | pCB1004-EV | / | / | / | pCB1004 | Empty vector | Yoshida et al., 2009 |
| Rice stable transgenics | pSC038 | / | / | / | pCambia1300 | pRGA4::RGA4genomic | Cesari et al., 2013 |
| | pSC086 | / | / | / | pCambia2300 | pRGA5::RGA5genomic | Cesari et al., 2013 |
| | pADu52.3 | Quikchange | pAHC17-pRGA5::RGA5 | oSC385/oSC386 | pCambia2300 | pRGA5::RGA5m1 | Okuyama et al., 2011 |
| | pADu50.3 | Quikchange | pAHC17-pRGA5::RGA5 | oSC332/oSC333 | pCambia2300 | pRGA5::RGA5m2 | Okuyama et al., 2011 |
| | pADu51.3 | Quikchange | pADu52.3 | oSC332/oSC333 | pCambia2300 | pRGA5::RGA5m1m2 | |
| SPR | pCV64 | / | / | / | pET15b | dSP-AVR-Pia | de Guillen et al., 2015 |
| | pCV61.2 | / | / | / | pET15b | dSP-AVR1-CO39 | de Guillen et al., 2015 |
| | pLM004 | in fusion | synthetic gene | / | pDB_ccdb_his_3C | dSP-AVR-PikD | |
| | pLM003 | in fusion | synthetic gene | / | pDb_ccdb_pepL_his_3C | dSP-AVR1-CO39 T41G | |
| | pCV147 | recombination | pSC60 | oTK439/oTK334 | pET15b | dSP-AVR-Pia F24S | Ortiz et al., 2017 |
| | pLM012 | in fusion | synthetic gene | / | pDB_ccdb_his_MBP_3C | MBP:RGA5_HMA | |
| | pLM015 | in fusion | synthetic gene | / | pDB_ccdb_his_MBP_3C | MBP:RGA5_HMAm1 | |
| | pLM018 | in fusion | synthetic gene | / | pDB_ccdb_his_MBP_3C | MBP:RGA5_HMAm1m2 | |
| | pLM009 | in fusion | synthetic gene | / | pDB_ccdb_his_MBP_3C | MBP:Pikp-1_HMA | |
